# Disease-associated loci share properties with response eQTLs under common environmental exposures

**DOI:** 10.1101/2025.04.30.651602

**Authors:** Wenhe Lin, Mingyuan Li, Olivia Allen, Jonathan Burnett, Joshua M Popp, Matthew Stephens, Alexis Battle, Yoav Gilad

**Author notes:** Correspondence should be addressed to W.L., A.B. and Y.G. These authors contributed equally to this work (Wenhe Lin, Mingyuan Li).

## Abstract

Many of the genetic loci associated with disease are expected to have context-dependent regulatory effects that are underrepresented in the transcriptomes of healthy, steady-state adult tissues. To understand gene regulation across diverse environmental conditions and cellular contexts, we treated a broad array of human cell types with three environmental exposures *in vitro*. With single-cell RNA-sequencing data from 1.4 million cells across 51 individuals, we identified hundreds of response expression quantitative loci (eQTLs) that are associated with inter-individual differences in regulatory changes following treatment with nicotine, caffeine, or ethanol in diverse cell types. We also identified dynamic regulatory effects that vary across differentiation trajectories in response to exposure. In contrast to steady-state eQTLs, and similar to disease risk loci, response eQTLs are enriched in distal enhancers and are regulating genes that experienced strong selective constraint, contain complex regulatory landscapes, and display diverse biological functions. We identified response eQTLs that coincide with disease-associated loci not explained by steady-state eQTLs. Our results highlight the complexity of genetic regulatory effects and suggest that our ability to interpret disease-associated loci will benefit from the pursuit of studies of gene-by-environment interactions in diverse biological contexts.

## Introduction

Complex traits are modulated by a combination of genetic and environmental factors, where genetic effects act in a context-dependent manner^1^. Contexts are multi-dimensional and encompass not just spatial components, like the cell- or tissue-specific milieu; but temporal components, such as age; and external components, including exposures to a variety of environmental conditions. Genetic effects on gene regulation have been also shown to vary across different biological contexts, including tissue types^2^, cell states^3^, developmental stages^4^, and external perturbations^5^. Since dysregulation of gene expression in specific contexts can contribute to disease risk and progression, understanding how genetic effects operate across diverse conditions and contexts is essential. To do so, we need to expand the scope of eQTL studies to account for gene-by-environment (GxE) interactions and study gene regulation in multi-dimensional contexts. We need to understand not just where, but when, and under what circumstances critical, disease-relevant regulatory events are taking place.

However, studying gene regulation in humans *in vivo* presents significant challenges, particularly when investigating GxE interactions. A major hurdle lies in the sheer complexity of the problem: there are potentially millions of relevant environmental exposures, ranging from diet, pathogens, and toxins to stress and lifestyle factors, and thousands of distinct cellular contexts, each with unique regulatory landscapes influenced by cell type, developmental stage, and physiological state. Conceptually, this creates a vast, multi-dimensional matrix of GxE interactions, where each combination of cellular context and environmental exposure represents a coordinate that may influence gene regulation and, ultimately, disease risk. To fully understand how GxE interactions shape complex traits and disease, we need to systematically map this matrix. Only then can we begin to unravel how specific environmental factors impact gene regulation across diverse cellular contexts. Current experimental approaches, whether in model organisms or human *in vitro* cell-based systems, only scratch the surface. Animal models allow for controlled *in vivo* studies and offer powerful tools for mechanistic insights, but their applicability to human biology is limited, especially when considering the role of human-specific genetic variation that was shaped by unique demographic histories. In contrast, *in vitro* models using human cells provide a direct window into human gene regulation and are more amenable to controlled perturbation studies. However, *in vitro* studies have focused on a limited set of exposures and cell types, quite often using immune cells^6–16^ , leaving significant gaps in our understanding of how environmental factors impact gene regulation across many other disease-relevant cellular contexts.

For many environmental exposures, we lack fundamental knowledge about which cell types are most responsive or critical for mediating their effects. This adds another layer of complexity, as it is often unclear where to focus experimental efforts. Compounding this challenge is the fact that, with current methods, each dynamic eQTL study typically investigates a single environmental perturbation in a single cellular context^5,6,17,18^. While these studies provide valuable insights, the pace of data generation is too slow to keep up with the scale of the problem. Machine learning and AI models may eventually enable us to predict the effects of new environmental contexts. However, the success of such models will depend on the availability of high-quality, diverse datasets for training and evaluation. At present, we are far from achieving this goal.

To accelerate progress in mapping GxE interactions, there is a need for scalable approaches that can efficiently profile the effects of environmental exposures across a wide range of cell types. In this study, we leveraged heterogeneous differentiating cultures (HDCs) derived from human induced pluripotent stem cells (iPSCs) as a flexible and scalable system for investigating GxE interactions across diverse cellular contexts. HDCs offer a unique advantage: as iPSCs differentiate, they give rise to a complex mixture of cell types spanning all three germ layers, creating a dynamic and diverse cellular environment^19^. This diversity allows us to capture cell type–specific responses to environmental treatments without the need to isolate or engineer each cell type individually. While each experiment still focuses on a single environmental exposure at a time, the HDC system enables us to observe GxE effects across many distinct cell types simultaneously, significantly expanding the scope of GxE mapping compared to standard experimental approaches.

To demonstrate the utility of this strategy, we treated HDCs from 51 genetically diverse individuals with chemicals corresponding to three common, disease-relevant environmental exposures: ethanol, caffeine, and nicotine. Our experimental design enabled us to systematically profile GxE interactions across a broad spectrum of cell types for each exposure, capturing both shared and context-specific genetic effects. With this multi-context framework, we then explored the underlying mechanisms driving context-dependent gene regulation. We identified hundreds of treatment-specific and temporally dynamic eQTLs that would be missed in steady-state or single-cell-type studies. By characterizing the genetic architecture of these context-responsive loci, we uncovered how genetic variation, cell type identity, and environmental factors converge to shape gene expression. We further examined response eQTL overlap with disease-associated loci, shedding light on how GxE interactions contribute to disease risk.

## Results

We generated HDCs from the iPSCs of 51 unrelated Yoruba individuals from Ibadan, Nigeria (YRI). Starting on Day 2 after HDC formation, we subjected HDCs to chronic, repetitive treatment with either nicotine, caffeine, ethanol, or vehicle control (see Methods; **Fig1A**). On Day 21, we dissociated HDCs and multiplexed samples from each individual and condition in equal proportions in preparation for single-cell RNA-sequencing using the 10X platform (see Methods). We recorded extensive metadata at every stage of sample collection and processing (**TableS1**). Following rigorous quality control and filtering, we retained data from 1,416,512 high-quality cells from 191 HDC samples (**FigS1**) yielding a median of 5799 cells per sample and 6447 unique molecular identifier (UMI) counts per cell.

**Figure 1.**
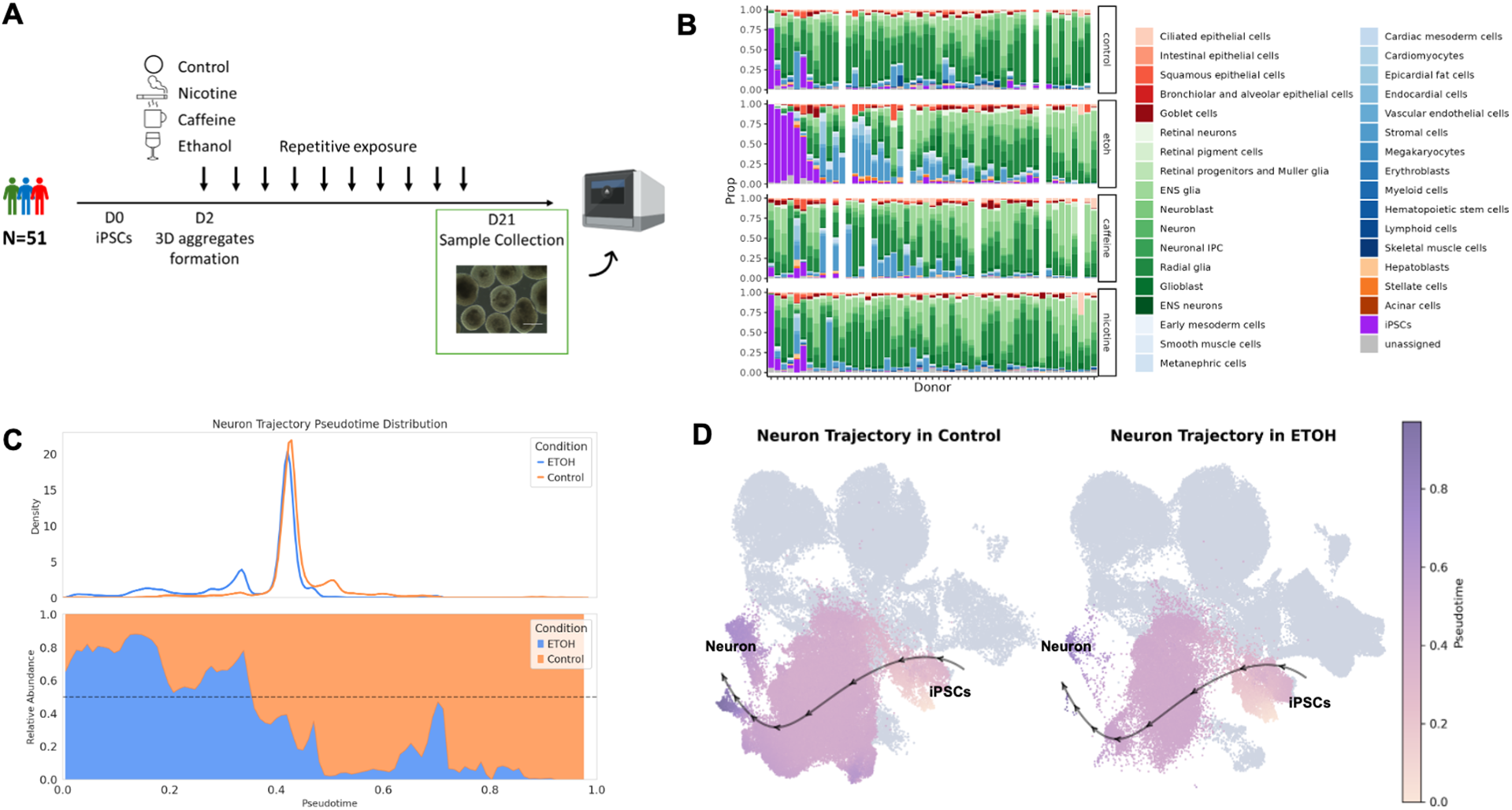
Overview of the study design and dataset. (**A**) A panel of 51 YRI iPSC lines were formed into 3D aggregates and differentiated simultaneously as HDCs. HDCs were separately exposed to nicotine, caffeine, and ethanol starting Day 2. On Day 21, scRNA-seq data were collected from control HDCs and treated HDCs. (**B**) Cell type proportion in HDCs across individual lines and conditions. HDCs include cell types from epithelium (red), ectoderm (green), mesoderm (blue), endoderm (orange), undifferentiated iPSCs (purple), and unassigned cells (gray). (**C**) Pseudotime distribution (top) and relative cell proportion in control versus ethanol (bottom) in the neuronal differentiation trajectory. (**D**) Visualization of the differentiation trajectory from iPSCs to neurons in control versus ethanol, shown in a shared UMAP embedding. Cells not assigned to the neuronal trajectory are shown in gray.

### Treatment-dependent effects on cell type composition and cell states

We started exploring the effects of each treatment by characterizing cellular heterogeneity within control and treated HDCs. To do so, we first compiled a reference list of cell type markers based on two previous large single-cell RNA sequencing datasets: the human fetal cell atlas from Cao *et al.* and the first-trimester fetal brain atlas from Braun *et al.* (**TableS2**). Using this curated reference list and a clustering-free multivariate statistical method^20^, we annotated a total of 1,394,531 (98.4%) cells to 34 cell types (Methods and **FigS2**). We used multiple approaches to evaluate our annotation results to indicate that our approach is robust and that the annotations are largely consistent with unsupervised clustering (Methods and **FigS3**).

We observed substantial inter-individual variation in cell type composition among control HDCs (**Fig1B**). Most lines were dominated by ectodermal cell types, consistent with previous studies, which have reported that iPSCs are prone to differentiating into neuronal cells^21–23^, even in directed differentiations for non-neuronal lineages^24^. We also observed clear differences in cellular composition across treatment conditions, indicating treatment effects on cellular differentiation and abundance (**Fig1B**).

We further examined the effects of treatment on cell state. As HDCs contain a complex mix of cell states, we reasoned that grouping cells into broad, mutually exclusive cell types may have limited our ability to detect subtle treatment-induced changes in cell abundance. Thus, we modelled cell states as overlapping neighborhoods on a K-nearest neighbors (KNN) graph^25^. Rather than assigning cells to predefined clusters, this approach allows cells to have partial membership in multiple ‘neighborhoods’ that represent fine-grained latent cell states. Considering each treatment independently, we tested for differential abundance of cells from control and treated HDCs in each neighborhood. For example, to assess the effects of ethanol on cell state, we assigned control and ethanol-treated cells to 10,860 neighborhoods, 6,038 of which showed evidence of differential abundance between control and treated HDCs at a 5% false discovery rate (FDR) (**FigS4A**).

To interpret the results, we tested for enrichment of discrete cell types within each neighborhood. We observed that most neighborhoods enriched with cells from ethanol-treated HDCs are dominated by iPSCs and mesoderm-derived cell types. In contrast, neighborhoods enriched with cells from control HDCs are populated largely by neuronal cells (**FigS4B**), consistent with previous work showing that ethanol inhibits neuronal differentiation^26–28^. We repeated the same analyses for HDCs treated with caffeine and nicotine. We found that hepatoblasts, epicardial fat cells, vascular endothelial cells, and stromal cells are more abundant in response to caffeine, while iPSCs and neuronal cells are less abundant (FDR < 0.05; **FigS4C-D**). We did not observe a significant effect of nicotine on either cell type abundance or cell state (**FigS4E-F**).

Our HDC dataset captures the continuum from naive to terminally differentiated cell states along multiple lineages, offering an opportunity to study gene regulation as a dynamic process. To directly model these continuous changes in cell state, we performed trajectory inference using Palantir^29^, which reconstructs the progression from stem-cell-like to terminal cell states, and assigns each cell a continuous pseudotime reflecting its position along this path. Using this approach, we identified 7 differentiation trajectories, towards epicardial fat cells, hepatoblasts, retinal neurons, neurons, vascular endothelial cells, ENS glias, and retinal pigment cells (**FigS5A** and **TableS3**). We assessed the inferred trajectories using marker genes expression. For example, hepatoblasts and epicardial fat cell trajectories share early and intermediate cells before the bifurcation of the mesoderm and endoderm differentiation trajectories (**FigS5B**), and their naïve, intermediate, and terminal state marker gene expression matches expected trends (**FigS5C**). Trajectory analysis of treated and control cells reinforces and expands upon our KNN-based analyses of cell state changes. For instance, ethanol-treated samples have a pronounced redistribution of cells along pseudotime in the neuronal trajectory, resulting in an enrichment of cells in earlier differentiation states and depletion of later-stage neuronal cells compared to controls (**Fig1C-D**), corroborating the KNN findings. Thus, trajectory analysis offers a dynamic view of treatment impact.

### Widespread transcriptional effects of the treatments

Next, we considered the effect of the treatments on gene expression within cell type. To characterize shared and treatment-specific effects on transcription, we performed differential expression analysis using the 25 HDC lines for which control and treated cells were cultured in parallel (see Methods). Using data from 25 cell types we annotated with high confidence (**FigS6**), we identified 10,023, 6,789, and 1,738 differentially expressed genes (DEGs) between control cells and cells treated with either ethanol, caffeine, or nicotine, respectively (FDR < 0.05; **TableS4**). The proportion of DEGs varies from 0.03% to 39% across cell types (**FigS7**), indicating that the different treatments (environmental exposures) have distinct regulatory effects in different cellular contexts. We found that DEGs are engaged in a variety of biological processes, with variation across cell types and treatments (**FigS8A**). Notably, genes known to be involved in the cellular response to toxic substances are strongly activated in hepatoblasts in response to all three treatments (normalized enrichment score (NES) = 1.8 to 2.6; **FigS8B**). We also observed transcriptional responses to treatment that recapitulate findings from previous studies. For example, in iPSCs, ethanol induces a transcriptional network enriched in cell proliferation genes (NES = 2.3, FDR < 4×10^-11^). Hepatic differentiation genes, such as *AFP* and *HNF4A*, are highly and specifically upregulated in response to caffeine in hepatoblasts (*AFP*: (logFC) = 3.1, FDR < 0.007; *HNF4A*: logFC = 1.6, FDR < 0.004), consistent with prenatal effects of caffeine in rodents^30^. Finally, genes relevant to fibrosis and apoptosis were activated in cardiomyocytes in response to nicotine (NES = 2.2, FDR< 3.7×10^-5^), reiterating findings from previous *in vitro* studies^31,32^.

### Regulatory effects of eQTLs are responsive to treatments

Having characterized global transcriptional responses to each of the three treatments, we sought to explore how treatment-induced transcriptional changes are shaped by the interaction between genotype and environment (the treatments). To identify genetic effects on gene expression (eQTLs), including both treatment-dependent eQTLs (response eQTLs) and standard (i.e., non-response) eQTLs, we considered data from a subset of 14 cell types that are represented by at least 30 individuals (see Methods; **FigS2**). First, we used a linear model to map *cis* eQTLs (MAF > 5%) in each cell type independently, performing a separate analysis for each treatment and control group (see Methods). We then jointly analyzed the effects of all top variant–gene pairs (n=9,734) identified in the control and treatment groups across all 14 cell types. By leveraging information across cell types, we discovered between 754 to 1,457 standard eQTLs for each treatment and cell type combination (local false sign rate (LFSR) < 0.1), corresponding to 6,198 eGenes across the entire dataset (**FigS9**).

To characterize response eQTLs in each cell type, we compared posterior genotype effect size estimates between control and treatment conditions. We defined response eQTLs as those for which the change in eQTL effect size between control and treatment groups differs by at least 1.5 fold. We identified between 31 to 110 response eQTLs in each cell type (LFSR < 0.1; **Fig2A**), corresponding to a total (combined across cell types) of 380, 336, and 276 eGenes in the ethanol, caffeine, and nicotine conditions, respectively (**TableS5**).

**Figure 2.**
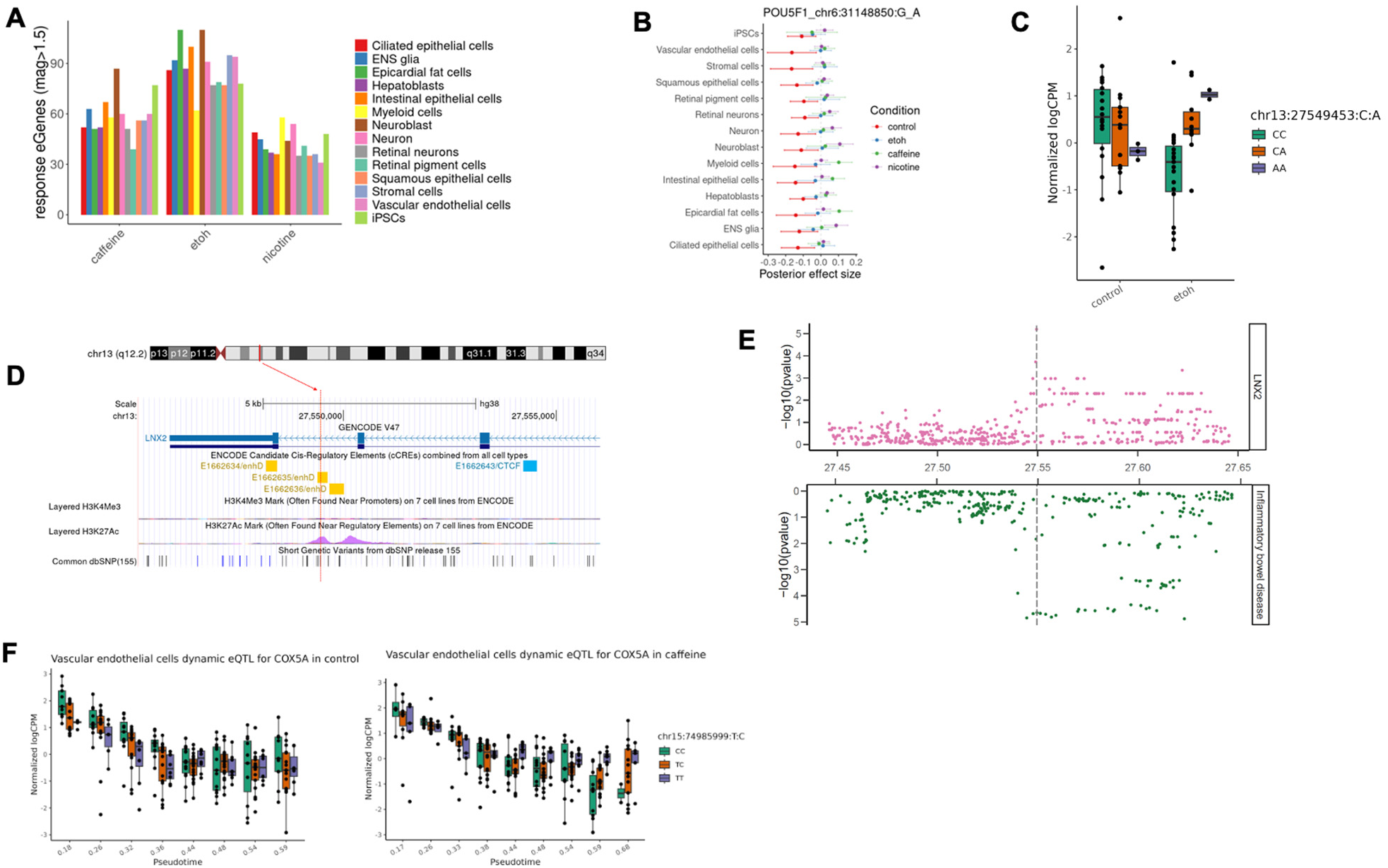
Regulatory effects in response to treatments. (**A**) Discovery of response eQTL-gene pairs in each treatment across 14 cell types. (**B**) A shared response eQTL (chr6:31148850:G_A) for *POU5F1*. The forest plot shows effect estimates of the eQTL across 14 cell types in control (red), ethanol (blue), caffeine (green), and nicotine (purple). Dots are posterior means, and error bars depict ±1 posterior standard deviations. (**C**) An ethanol-response eQTL (chr13:27549453:C_A, P_ethanol_=6.4×10^-6^) for *LNX2*. The eQTL effect emerged in response to ethanol and was present only in intestinal epithelial cells. (**D**) Genomic annotations of the ethanol-response eQTL for *LNX2* (chr13:27549453:C_A). The SNP is located on chromosome 13 and falls in an enhancer element (yellow) annotated by ENCODE, featured by a peak of H3K27Ac and a low activity of H3K4Me3. (**E**) Colocalization of the ethanol-response eQTL for *LNX2* (chr13:27549453:C_A) in intestinal epithelial cells with inflammatory bowel disease. The Manhattan plot shows significance levels of variants tested for association with gene expression level (top) or with the trait (bottom). Vertical lines depict the genomic location of the candidate colocalized variant. (**F**) A caffeine dynamic response eQTL (chr15:74985999:T_C, FDR = 0.065) for *COX5A*. The eQTL effect emerged in response to caffeine and was present only in late stages of vascular endothelial cell trajectory.

Response eQTLs include both shared and cell-type-specific effects. For example, a response eQTL for *POU5F1* (*OCT4*), which encodes a transcription factor critical for maintaining stem cell pluripotency, is widespread among untreated cell types, but its effect is consistently diminished in all treated conditions (**Fig2B**). In myeloid and epicardial fat cells, however, the direction of the eQTL effect is reversed in response to caffeine. We also observed response eQTLs unique to one cell type. For example, an eQTL located in an enhancer element for *LNX2,* emerged in response to ethanol and is present only in intestinal epithelial cells (**Fig2C, 2D**). Moreover, it colocalizes with a candidate GWAS locus associated with inflammatory bowel disease (IBD; posterior probability of sharing the same region (PP_regional) = 0.92; **Fig2E**), suggesting a functional mechanistic connection between a gene regulatory variant, ethanol exposure, *LNX2* expression in the intestinal epithelium, and IBD. Notably, this ethanol-dependent eQTL was not identified as an eQTL in any of the 54 steady-state tissues included in GTEx^2^

### Dynamic genetic regulation along differentiation trajectories

We also leveraged the inferred pseudotime from the trajectory analysis to systematically identify dynamic eQTLs whose genetic effects vary across differentiation states captured by pseudotime. To achieve this, we evaluate a genotype-by-pseudotime interaction term in a linear model, following our previous dynamic eQTL mapping approach^19,33^. We applied a genome-wide FDR cutoff of 0.1 using EigenMT adjusted p-values for the pseudotime interaction effect. We identified 5,119 dynamic eQTLs corresponding to 614 unique eGenes across treatments and trajectories (**TableS6**).

To evaluate the robustness of our approach we performed replication analysis focusing on neuronal trajectory dynamic eQTLs identified under control conditions. We compared these results with a previous dataset that examined neural differentiation in HDCs^19^. A majority of our identified dynamic eQTLs were replicated in this previous study (π₁ = 0.77; **FigS10A-B**). We further explored the temporal nature of these dynamic eQTL effects, categorizing them as early (21%) or late (79%) based on their change of effect sizes throughout the differentiation trajectory. Of particular interest were the dynamic eQTLs whose effects appeared exclusively in treatment conditions (dynamic response eQTLs), indicating that environmental treatments can dynamically modulate genetic regulation during cellular differentiation. For instance, we identified a caffeine-response dynamic eQTL affecting *COX5A* (encoding a mitochondrial cytochrome c oxidase subunit) whose regulatory effect emerged only in late stages of vascular endothelial cell differentiation in the caffeine treatment and was not discovered in the mapping of discrete cell types (**Fig2F**). This dynamic response eQTL overlaps with a variant associated with mean corpuscular volume, as annotated by OpenTargets^34^, highlighting a potential functional connection between caffeine-induced changes over the course of vascular development and a hematological trait.

### Response eQTLs share genomic features with disease-associated loci

Many disease-associated loci in putatively regulatory regions of the genome are not annotated as steady-state eQTLs in standard (steady-state) assays, including those identified using the GTEx Project data^2^. It has been argued that standard eQTLs and disease-associated loci systematically differ with respect to genomic location, selective constraint, regulatory complexity, and functional enrichments^35^. To determine whether response eQTLs are distinct from standard steady-state eQTLs, we compared locus-level and gene-level features of response eQTLs and standard eQTLs identified in HDCs.

When we separately examined response eQTLs and standard eQTLs, a distinct localization pattern emerged, with strong enrichment of standard eQTLs near TSS, while response eQTLs showing only a modest, much less pronounced enrichment near TSS (**Fig3A**). This pattern is largely consistent across all 14 cell types and treatments (**FigS11**), suggesting that steady-state eQTLs and response eQTLs may perform different regulatory roles.

**Figure 3.**
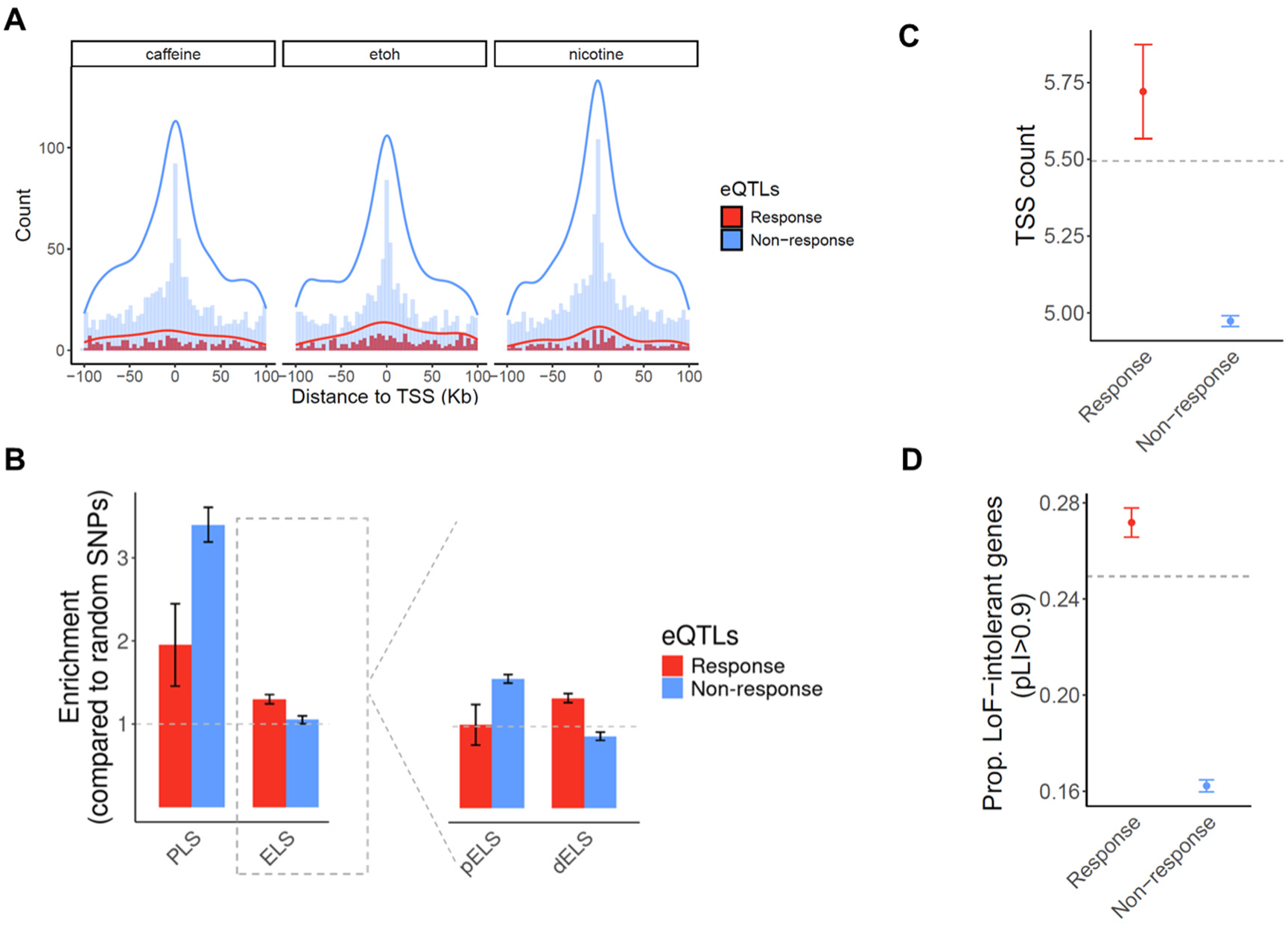
Properties of response eQTLs/eGenes and non-response eQTLs/eGenes. (**A**) Distance of response eQTLs (red) and non-response eQTLs (blue) to the TSS of the target eGene. The overlaid histograms show distance-to-TSS distribution in each group of eQTLs in 5Kb bins. The curves show the density estimate for each group of eQTLs. (**B**) Enrichment of response eQTLs (red) and non-response eQTLs (blue) in promoter-like signatures (PLS) and enhancer-like signatures (ELS). The right plot compares enrichments of response eQTLs and non-response eQTLs in proximal enhancers (pELS) and distal enhancers (dELS). Enrichment level is expressed as the odds ratio from Fisher’s exact test. (**C-D**) Genic features of response eGenes (red) and non-response eGenes (blue). Horizontal dashed lines depict the average number of the corresponding feature across all genes that were tested in eQTL mapping. (**C**)Average number of TSS. (**D**) Fraction of selectively constrained genes (pLI > 0.9). Error bars represent standard deviations.

We further characterized the eQTLs found in our study using the annotation of *cis* regulatory elements (CREs) from ENCODE^36^. We found that both standard and response eQTLs under each treatment are enriched in promoter regions, with steady-state eQTLs showing a markedly stronger enrichment (OR_average_=3.4, P_combined_=3.63×10^-38^; **Fig3B**). Response eQTLs are also enriched in enhancers (OR_average_=1.29, P_combined_=2×10^-3^) but steady-state eQTLs are not. This was intriguing, so we explored this further by partitioning enhancers to proximal (within 2kb of a TSS) and distal (beyond 2kb from a TSS) with respect to the gene they regulate. We found that steady-state eQTLs are enriched in proximal enhancers (OR_average_=1.58, P_combined_=2.8×10^-12^) and depleted in distal enhancers (OR_average_=0.88, P_combined_= 0.01). Response eQTLs show the opposite pattern: they are enriched in distal enhancers (OR_average_=1.35, P_combined_=1×10^-3^), but are neither enriched nor depleted in proximal enhancers (OR_average_=1.02, P_combined_=0.7).

To further investigate the genetic architecture of treatment-responsive gene expression, we analyzed the target genes that are associated with response eQTLs (response eGenes) and compared them to genes associated with steady-state eQTLs (non-response eGenes). For each group of eGenes, we assessed multiple genic features, including regulatory complexity, evidence of selective constraint, and functional enrichment.

Regulatory complexity can be achieved in a number of ways; for example, by differential usage of TSS across cell types^37,38^, or by long enhancer sequences that contain additional binding sites for transcription factors^39–41^. To determine whether response eGenes are associated with increased TSS usage, we used publicly available summary statistics from the FANTOM project to compute the number of TSS used by each gene across 975 human samples^38^. Across all genes, the average number of TSS is 5.49, but that non-response eGenes have significantly fewer TSS (4.79 TSS on average, compared to 5.73 TSS for response eGenes; P=0.001; **Fig3C**). To assess enhancer activity associated with eGenes, we calculated the cumulative enhancer length for each eGene across 131 tissue or cell types using enhancer-gene predictions from an activity-by-contact model^42^. Our analysis revealed that response eGenes have a slightly longer cumulative enhancer length per active tissue/cell type compared to non-response eGenes (P=0.06; **FigS12**). These results suggest that response eGenes may have a greater level of regulatory complexity.

Previous studies indicated that selectively constrained genes are depleted for steady-state eGenes^43^. Consistent with this, we found that genes depleted of loss-of-function (LoF) variants (as measured by the pLI score; see Methods) are underrepresented among non-response eGenes (**Fig3D**). In contrast, response eGenes show an enrichment of LoF-intolerant genes, with a significantly higher proportion of high-pLI genes compared to non-response eGenes (P=8.6×10^-6^; **Fig3D**). Although stronger selective pressure is often associated with lower minor allele frequency (MAF), we observed that response eGenes tend to have higher MAF than non-response eGenes (**FigS13**). These observations indicate that eGenes associated with context-dependent regulatory effects may experience different selective pressures than steady-state, non-response eGenes.

Previous studies have reported that while disease-associated genes are enriched across a wide range of functional categories, steady-state eQTLs are often depleted in these same categories^35^. In our study, we found that response eGenes show enrichment in similar functional categories as disease-associated genes and confirmed that steady-state eQTLs are indeed depleted in these categories (**FigS14A**). Although the enrichment of response eGenes varies across cell types and treatments, we consistently observed that response eGenes are comparatively more enriched in most functional categories than non-response eGenes (**FigS14B**).

### Response eQTLs provide insight into the role of GxE in pathology

To investigate whether response eQTLs offer functional insight into disease mechanisms, we integrated our data with GWAS results to evaluate colocalization of eQTLs and disease-associated loci. We obtained GWAS summary statistics for 265 traits, including 220 human phenotypes from a cross-population atlas^44^ and 45 blood or immune-related traits from Fair *et al*^45^. Using a Bayesian colocalization method^46^, we identified hundreds of regions where eQTLs colocalized with at least one trait (PP_regional > 0.5; **TableS7**). We found that the proportion of colocalized loci is universally higher for response eQTLs (an average of 4.8%) than for non-response eQTLs (an average of 0.4%), regardless of cell type or treatment (**Fig4A**). This suggests that in the context of our GxE experiment, response eQTLs are more likely to be functionally relevant to complex traits. This may be especially true when response eQTLs are active in a disease-relevant cellular context.

**Figure 4.**
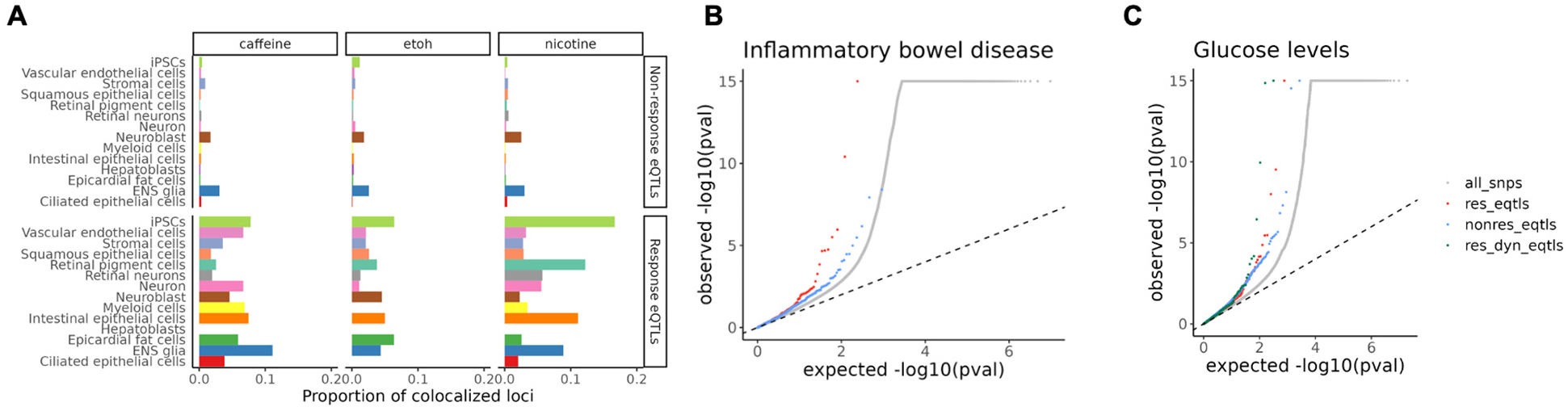
Enrichment of response eQTLs in GWAS signals for complex traits. (**A**) Proportion of non-response eQTLs and response eQTLs colocalizing with GWAS hits for any trait for each treatment, colored by cell type. (**B**) QQ plot of GWAS signals for inflammatory bowel disease, grouped by categories of SNPs. The grey dots are all SNPs, red dots are response eQTLs, and blue dots are non-response eQTLs. (**C**) QQ plot of GWAS signals for glucose levels, grouped by categories of SNPs. The green dots are dynamic response QTLs.

For example, we identified colocalization of caffeine response eQTLs with trait-associated loci in ENS glia and intestinal epithelial cells, some of which are known to be linked to gastrointestinal diseases. Specifically, a caffeine response eQTL for *SERBP1* identified in intestinal epithelial cells shows a strong colocalization with a GWAS variant for IBD (PP_regional=0.88; **FigS15**).

It is established that disease-associated loci are typically enriched among regulatory QTLs, including eQTLs. To explore this in our data, we compared the enrichment of GWAS signals in non-response eQTLs, response eQTLs, and dynamic response eQTLs (identified using the trajectory analysis), for the 265 complex traits. We found that incorporating these additional layers of context-specificity enabled us to identify regulatory effects that would have been missed otherwise. For example, using GWAS summary statistics for IBD, we found a stronger enrichment of low GWAS *P*-values in caffeine-response eQTLs compared to that of non-response eQTLs (**Fig4B**). Additionally, dynamic response eQTLs, showed even higher enrichment for certain traits, such as glucose levels, compared to both non-response and response eQTLs (**Fig4C**). For many other traits, response and dynamic eQTLs capture disease relevant genetic associations that were missed by steady-state eQTLs (**FigS16, S17**). Together, our results suggest that regulatory effects in response to environmental exposures and across differentiation in diverse cell types can improve our understanding of disease-associated loci.

## Discussion

Although GWAS signals are broadly enriched for eQTLs^47^, steady-state eQTLs have failed to explain an appreciable fraction of disease-associated loci. A recent analysis by Mostafavi et al. found systematic differences between the features of steady-state eQTLs and disease-associated loci with respect to their genomic location, regulatory complexity, selective constraint, and functional enrichment (**Table 1**). Mostafavi et al. findings suggested that context-specific eQTLs may provide additional mechanistic explanations for disease-associated loci that are putatively involved in gene regulation. The challenge is that the experimental setup required to identify eQTLs in just a single specific context (such as a single combination of cell type and treatment exposure) is resource intensive, requiring adequate sample size and careful control of experimental conditions, making it difficult to scale across many contexts.

**Table 1.** Summary of characteristics for GWAS hits, steady-state eQTLs, and response eQTLs. *The table is adapted from Mostafavi *et al*^35^.

|  | Feature | GWAS hits | Steady-state eQTLs | Response eQTLs |
| --- | --- | --- | --- | --- |
| SNP-level | Distance to TSS | Distal | Proximal | Distal |
|  | Promoter | Less enriched | More enriched | Less enriched |
|  | Enhancer | Distally enriched | Proximally enriched | Distally enriched |
| Gene-level | Selective constraint | High | Low | High |
|  | Regulatory complexity | High | Low | High |
|  | Biological function | Enriched | Depleted | Enriched |

Our work highlights the utility of HDCs, a scalable and efficient human *in vitro* model for studying GxE interactions across differentiation stages in diverse cell types. While our system still only allows us to examine the effect of one environmental exposure at a time, the ability to profile its effects across dozens of diverse cell types and states represents a significant advance. Beyond improving data generation efficiency, data from HDCs may ultimately provide a foundation for predictive models that can extend across the broader landscape of GxE interactions, helping to unravel the regulatory architecture of complex traits and diseases.

The use of HDCs allowed us to uncover hundreds of gene regulatory variants with dynamic effects, many of which have never been observed in steady-state adult tissues. For example, we identified an ethanol response eQTL for *LNX2* in intestinal epithelial cells that colocalized with a GWAS variant for IBD^48^. This eQTL was not among those detected in steady-state GTEx tissues, despite a much larger sample size^2^. Additionally, using dynamic eQTL mapping, we identified genetic variants with regulatory impacts that change across developmental trajectory pseudotime in response to environmental treatments. For instance, we uncovered a caffeine-responsive dynamic eQTL for *COX5A* in vascular endothelial cells differentiation. Had we only explored a single cell type or failed to account for GxE and temporal interactions, these and many other context-dependent, functionally informative eQTLs would not have been detected.

Comparing response eQTLs to standard eQTLs, we found that the two types of loci are associated with distinct regulatory architectures and functional properties. For example, unlike steady-state eQTLs, response eQTLs are enriched near distal regulatory elements, which have frequently been associated with context-specific gene regulatory activity^49–51^. This suggests that response eQTLs may function through long-range regulatory interactions, possibly integrating environmental signals to modulate gene expression in a context-dependent manner. In addition, response eGenes have greater regulatory complexity than steady-state eGenes, showing an increase in differential TSS usage and cumulative enhancer length – both of which suggest potential for fine-tuning gene expression levels and generating transcriptional diversity. Response eGenes are also more likely to be functionally important and subject to selective constraint, suggesting that their cognate eQTLs may help to explain inter-individual differences in disease risk.

Our analysis revealed a strong resemblance between response eQTLs and GWAS loci, which share a number of SNP-level and gene-level features in common (**Table 1**). In the context our study, we also found that response eQTLs are more likely to colocalize with disease-associated variants than steady-state eQTLs. This suggests that context-dependent response eQTLs, which are underrepresented in steady-state tissues, play a critical role in mediating the effects of non-coding disease-associated loci. Our findings are consistent with observations from a recent study of gene-diet interactions in baboons, where we identified distinct patterns separating standard eQTLs from diet-response eQTLs^52^. In addition to showing a significant enrichment among GWAS signals for metabolic traits, diet-response eQTLs exhibit complex regulatory architectures resembling those observed for response eQTLs in the current study.

Unlike standard eQTLs, response eQTLs from both the current and the baboon study share similar characteristics with disease-associated loci. The concordance between these two models – one in live baboons and one in cultured human cells – suggests that context-dependent response eQTLs are likely to enhance our understanding of complex traits in general, whether they emerge in response to diet, chemical exposures, or other perturbations. In particular, we identify GWAS loci that do not overlap with steady-state eQTLs but are explained by response and dynamic eQTLs. We expect that by continuing to expand our focus beyond steady-state tissues to include a wider range of cell types, time points, and environmental exposures, we will uncover more context-dependent and functionally relevant regulatory loci, thereby enhancing our ability to explain inter-individual differences in disease risk.

### Limitations

While our study provides valuable insights into the mechanisms of GxE interactions, ethanol, caffeine, and nicotine are only a few of the myriad environmental factors that influence human health. Future studies should expand this approach to include a broader range of perturbations, such as dietary components, pollutants, and pathogens. In addition, larger datasets that include a wider range of genetic backgrounds and cell types would improve our ability to detect response eQTLs, particularly those with low allele frequencies, cell-type-specific effects, and small effect sizes.

#### Data and code availability

The single-cell RNA-seq data have been deposited at GEO under accession GSE292909. All original code, data, and summary statistics presented in this paper have been deposited at Zenodo (https://doi.org/10.5281/zenodo.15307006). Any additional information required to reanalyze the data reported in this paper is available from the lead contact upon request.

## Acknowledgments

We thank N. Gonzales for editing and providing comments on the manuscript; members of the Gilad lab and the Battle lab for valuable discussions; University of Chicago Research Computing Center for providing computational resources. This work was supported by NIH grant R35GM131726 to YG.

## Author contributions

Y.G. conceived the study; W.L. designed the study, performed all the experiments, and carried out the analysis with contributions from M.L.; O.A. and J.B. assisted W.L. with iPSC culture and HDC dissociation; M.L. performed trajectory inference and dynamic eQTL mapping; W.L. wrote the manuscript with contributions from M.L., Y.G., A.B., and J.P.; M.S. guided W.L. on mashr analysis; Y.G. and A.B. supervised the study.

## Methods

### Cell culture and differentiation

HDCs were formed using a modified version of the STEMCELL Agrewell 400 protocol. We cultured iPSCs in E8 media with rock and synched lines up in each batch to desired density (50%-90% confluency). On the seeding day, we first prepared Aggrewell 400 24-well plates by pre-treating wells with anti-adherence rinsing solution (07010, STEMCELL), followed by rinsing with warm basal media DMEM/F12. iPSCs were dissociated into single cells and resuspended in Aggrewell EB Formation Medium (05893, STEMCELL) with ROCK inhibitor Y-27632 and Penicillin/Streptomycin. We counted cells and recorded viability twice using Countess II. Cells were seeded into each prepared well at a density of 1,000 cells per microwell (1.2×10^6^ cells and 2mL per well). After 24 hours, we changed half of the media with fresh Aggrewell EB Formation Medium without ROCK inhibitor. After another 24 hours on day 2, we gently collected the aggregates and transferred them to an ultra-low attachment 6-well plate (CLS3471-24EA, Sigma) in prepared media for vehicle control or treatment groups. We maintained HDCs for an additional 19 days, replacing media with fresh prepared media every 48 hours.

For the chemical treatments, we used concentration of 80mM, 0.6mM, 10uM for ethanol (BP2818500, Fisher Scientific), caffeine (C0750-5G, Sigma), and nicotine (N3876-5ML, Sigma) treatment, respectively. We prepared 10,000X stock solutions in non-denatured 200 proof ethanol (BP2818500, Fisher Scientific) and stored aliquots in -20C for all batches. Each time of changing media, we prepared fresh media by adding an aliquot of corresponding substance in basal media (E6 media (A1516401, ThermoFisher Scientific) with Penicillin/Streptomycin). For vehicle control, we used fresh basal media with 0.01% 200proof ethanol. To avoid sample swaps, we separated samples by condition with HDCs exposed to the same treatment in the same 6-well plate where individual lines were separated by well. Each well contained HDCs from one line and each 6-well plate contained six lines of HDCs in the same condition. Culture plates for ethanol treatment group were placed in a dedicated incubator that contained an open dish at the bottom with a solution of 80mM ethanol to create an equivalent concentration of ethanol atmosphere and prevent ethanol evaporation in the media.

HDCs were dissociated on day 21. We harvested HDCs following the STEMCELL Agrewell400 protocol. After rinsing HDCs with phosphate-buffered saline (Corning 21-040-CV), we dissociated them with AccuMax (STEMCELL 7921) at 37C. We gently pipetted HDCs up and down for 30s every 5 min until HDCs were completely dissociated. We then quenched the dissociation with warm E6 media. After spinning down, cells were resuspended in cold E6 media and strained them through a 40 mm strainer (Fisherbrand 22-363-547). We counted cells and measured viability using Trypan Blue and Countess II (AMQAF1000, invitrogen). We then mixed cells from each line for the same condition together in equal proportions, with one mixed suspension per condition. Each mixed suspension was counted again using hemocytometer to determine the final cell concentration. We adjusted the final concentration to 800-1,200 cells/ul for each mixed suspension. Cell suspensions remained on ice before loading to the 10X Chromium Controller.

We split 51 iPSC lines into two batches. In the first batch (25 lines), we included both untreated control HDCs and HDCs from all three treatment groups, which we formed, maintained, and dissociated in two sub-batches. We processed all samples in the first batch in parallel for library preparation and sequencing. The second batch (26 lines) included treated HDCs from each treatment group but not controls. For additional controls, we used data from naïve HDCs that were generated from the same individuals in a previous study. We did this for two reasons. First, using the data from the controls included in the first batch, we found only minimal variation in gene expression between newly collected and previously collected HDC data. Second, we only planned to use data from both batches for cis eQTL mapping, an analysis that is not sensitive to technical batch effects (i.e., because genetic variation is unlikely to correlate with batch assignment). To assess differential expression and abundance between treated HDCs and controls, we only used samples processed in the first batch.

### Single-cell RNA sequencing

We generated scRNA-seq libraries using Chromium Next GEM Single Cell 3’ HT Reagent Kits v3.1 (1000348, 10X Genomics). Using the evenly pooled mix of lines from each condition in each sub-batch, we loaded a 10x chip targeting 30,000 cells per channel of the 10x chip and loaded the same pool of cells across multiple lanes to recover 6,000 cells per individual line.

After cDNA amplification and cleanup, samples from each sub-batch were stored at -20C. We processed all samples from sub-batches in parallel in each batch for library preparation and sequencing. Libraries were sequenced with a target depth of 30,000 reads per cell on the NovaSeq X in the University of Chicago Functional Genomics Core. We recycled the data from naïve HDCs of 26 individual lines that were generated from a previous collection^19^. These 26 lines are matched to the treated lines we generated in batch 2. We subset cells to 7,000 cells per line to align with our current collection.

We used CellRanger (7.0.0) to align samples to the human genome (GRCh38). We used Vireo to demultiplex samples by genotype and assign droplets to individuals^53^. We used a reference panel from 1000 Genomes Project Phase 3^54^ for demultiplexing. We removed doublets and droplets that could not be confidently assigned to an individual. We removed individuals with fewer than 50 assigned cells. We further filtered cells to keep only those with less than 20% mitochondrial reads and with at least 750 genes expressed and at most 9000 genes detected. After the quality controls, we obtained 1,416,512 high-quality cells for downstream analyses. For quality controls on the gene level, we used different filtering in different analyses based on the need and purpose, which are noted in each corresponding section.

### Cell type annotation

After extensive testing and evaluation, we chose to annotate cells at individual cell level using CelliD^20^ with some modifications based on our dataset. CelliD is a clustering-free multivariate statistical method that performs gene signature extraction and functional annotation for each individual cell in a single-cell RNA-seq dataset. It uses Multiple Correspondence Analysis (MCA) to produce a simultaneous representation of cells and genes in a low dimension space. Genes are then ranked by their distance to each individual cell, providing unbiased per-cell gene signatures. Then cells are annotated by evaluating the enrichment of per-cell gene signatures in a reference list of cell type markers.

To compile a reference marker list, we used two published datasets from the human fetal cell atlas^55^ and first-trimester brain atlas^56^. For the human fetal cell atlas, we first obtained the count matrix and filtered out low-quality cells. After removing placental cells, uncharacterized cell types, and rare cell types that have fewer than 100 cells, we obtained 56 cell types (156,430 cells). Next, we randomly subsampled 1,000 cells per cell type, filtered genes to protein-coding genes that are expressed in more than 10 cells, normalized the expression matrix by library size factors, log transformed the data, and selected highly variable genes. Then, we ran MCA and extracted top 250 genes for each cell type as their markers. We iterated this cell subsampling and preprocessing 20 times and used intersection as the final marker set for each cell type, with a median of 168 genes.

For the human first-trimester brain atlas, we used the same strategy. After quality controls, we first subset the data to a maximum of 20,000 cells per Celltype_TopLevelCluster (7 cell types, 57 Celltype_TopLevelClusters, 831,953 cells). Here, we kept a finer resolution of annotation as used in the original dataset to generate a more accurate reference marker list, and after cell annotation, we merged the annotated cells from the same cell type but different TopLevelClusters. We iteratively subsampled 1,000 cells per Celltype_TopLevelCluster and extracted top 250 genes using MCA. After 20 iterations, we used the intersection as the final marker set for each Celltype_TopLevelCluster, with a median of 157 genes.

To avoid redundancy, we excluded brain-derived cell types from fetal cell atlas: purkinje neurons, inhibitory neurons, astrocytes, granule neurons, oligodendrocytes, nhibitory interneurons, microglia, limbic system neurons, excitatory neurons, and unipolar brush cells. We combined reference marker lists generated from these two atlases and manually added a set of markers for iPSCs. Altogether, we obtained a final list of 114 cell types (clusters) as our reference marker list, with a total of 6470 genes.

To learn gene signatures of individual cells in our query dataset, we used a similar pipeline. After normalization and log transformation, we selected top 2,000 highly variable genes. Using the union of reference markers and the 2,000 highly variable genes, we performed MCA and extracted top 300 genes for each individual cell in the query dataset. To avoid the effects of cell cycling, we excluded ribosomal, mitochondrial, and cell-cycle genes in both reference markers and query gene signatures. Finally, we ran hypergeometric tests against the reference marker gene list to find the strongest enrichment of per-cell gene signatures among cell types (clusters) in the reference. Cells with a corrected p-value smaller than 0.01 are labelled as unassigned (20,772 out of 1,416,512). After assigning individual cells in the query dataset to reference cell types (clusters), we cleaned the annotations by combining similar cell types. First, we combined the labels from the same cell type, but different TopLevelClusters that are derived from the brain atlas. For cell types in the fetal atlas, we combined ENA glia and schwann cells; cardiomyocytes and cardiac progenitor cells; amacrine, horizontal, ganglion, photoreceptor, and bipolar cells as retinal neurons; ENS neurons, sympathoblasts, chromaffin, and visceral cells; epicardial fat and mesothelial cells; lymphatic and vascular endothelial cells; lymphoid and thymocyte; fibroblast, stromal, and mesangial cells; parietal and chief cells and goblet cells. After cell label cleaning, we have 1,416,512 cells assigned to 34 cell types and unassigned cells, with a median of 10,542 cells per cell type.

After removing unassigned cells and manually examining canonical marker gene expression, we identified 25 high-confidence cell types to use in downstream analyses (**FigS5**). These 25 cell types comprise a variety of diverse cell types spanning ectoderm, mesoderm, endoderm, and epithelium.

### Differential abundance analysis

We used milo to detect differences in cellular abundance without relying on cell type annotations^25^. Milo models cell states as overlapping neighborhoods on a KNN graph and then tests for differential abundance in each neighborhood. For each treatment, we first computed an KNN graph on the 25 lines of control and treatment group from batch 1 using 50 nearest-neighbors and 30 dimensions. Next, we defined neighborhoods on the KNN graph using a 5% randomly sampled cells. Then, we counted the number of cells from each sample and tested for differential abundance in each neighborhood. Milo implements this hypothesis testing in a Negative Binomial generalized linear model framework where we accounted for sex and batch effects. P-values were corrected using an adaptation of the Spatial FDR correction to account for the amount of overlap between neighborhoods. Specifically, each hypothesis test p-value is weighted by the reciprocal of the kth nearest neighbor distance.

### Differential expression analysis

We performed differential expression (DE) analysis between control and a treatment in each cell type using data from batch 1 (25 lines, 2 sub-batches) where we generated untreated and treated HDCs in parallel. We used dreamlet which applies pseudobulk approach and uses a linear mixed model with precision weights for single-cell RNA-seq datasets^57^. First, we removed samples (batch, cell type, condition, and individual combinations) with fewer than 10 cells, and removed cell types with fewer than 3 individuals. We kept genes with at least 10 cells with non-zero reads across all cells from control and one treatment. Next, we aggregated counts across cells from the same sample. We filtered samples and genes, normalized aggregated counts, and computed precision weights by number of cells using the function *processAssays.* In each cell type, we set treatment, sex, and batch as fixed effects and individual as a random effect in a linear mixed model. Finally, we fit the model on each gene using the Satterthwaite approximation for the hypothesis test by default and applied empirical Bayes shrinkage on the linear mixed model. Genes with an FDR-adjusted p-value < 0.05 were considered DE genes.

### Gene set enrichment analysis

We performed gene set enrichment analysis (GSEA) using the R package fgsea. fgsea is a powerful method that quickly estimates arbitrarily low GSEA p-values accurately based on an adaptive multi-level split Monte-Carlo scheme^58^. We used pre-ranked t-statistics for all tested annotated genes in each tissue from DE analysis as input and 50 hallmark gene sets from the Molecular Signature Database (MSigDB v7.5.1) as reference gene sets. The hallmark gene sets summarize and represent specific well-defined biological states or processes and display coherent expression^59^. These gene sets were generated by a computational methodology based on identifying overlaps between gene sets in other MSigDB collections and retaining genes whose gene expression levels are coordinate. Multiple testing was corrected using the Benjamini-Hochberg procedure and an FDR ≤ 0.05 and absolute NES > 1 was considered as significant enrichment. NES is a normalized enrichment score used to account for the size of a gene set and correlation between a gene set and genes in the ranked list.

### Trajectory isolation and pseudotime inference

We first applied contrastiveVI^60^, a deep generative model designed to deconvolve scRNA-seq data into shared and condition-specific latent spaces. To prepare the data, we filtered out cells with fewer than 1,000 unique molecular identifiers (UMIs) and excluded genes expressed in fewer than 10 cells. We then identified the top 2,000 highly variable genes (HVGs) using log-normalized counts and Scanpy’s *highly_variable_genes* with flavor *cell_ranger* for variable gene selection^61^.

Using contrastiveVI’s autoencoder framework, we trained a model on the raw UMI count matrix of the 2000 HVGs to extract 30 shared and 30 treatment-specific latent variables. The shared latent variables, which represent biological variation common across treatment and control conditions, were used as input for downstream trajectory analysis. To visualize the dataset, we constructed a neighborhood graph from the shared latent space and applied UMAP using Scanpy’s default parameters. This 2D embedding was used solely for visualization and did not influence trajectory inference or any QTL analyses.

Next, we used Palantir for trajectory inference^29^. To address computational burden, we first randomly selected 100,000 single cells regardless of cell types, and individuals in each condition. Then, we computed 30 diffusion components (DCs) from the neighborhood graph on these 400,000 cells using *palantir.utils.run_diffusion_maps*, which projects the data into a lower-dimensional space while preserving the cell-cell similarity structure. Terminal states were selected based on the extrema of DCs in terminally differentiated cell types. We then ran *palantir.core.run_palantir* with 30,000 waypoints to reconstruct differentiation trajectories, with the pseudotime ordering and trajectory probabilities toward each terminal states as outputs. Next, we used KNN regression with 30 nearest neighbors to map the rest of single cells with pseudotime and trajectory probabilities.

### Molecular cis-eQTL mapping

cis-eQTL mapping was performed using a linear regression model implemented by MatrixEQTL^62^. Genotypes were obtained from high-coverage (30x) whole-genome sequencing data from 1000 Genomes Project. We filtered SNPs in each cell type and condition combination by allele frequency and genotype frequency. We removed outlier SNPs with MAF < 0.05. We also filtered out loci with < 2 homozygotes across samples in each cell type and condition combination.

First, we removed samples (batch, cell type, condition, and individual combinations) with fewer than 3 cells, and removed cell types with fewer than 10 individuals. We kept genes with at least 5 cells with non-zero reads in each condition. Next, we aggregated counts across cells from the same sample. After count aggregation, we removed samples with no cells and genes with fewer than 2 donors with at least 3 aggregated reads. We used lenient filtering to include more genes to have enough commonly expressed genes when jointly analyzing data across cell types and conditions in mash analysis. In each cell type, we normalized the aggregated counts of all pseudobulk samples across control and the three treatment groups for a given gene using trimmed mean of M-values (TMM). Log-transformed normalized reads for each gene were inverse normal transformed across samples. For eQTL mapping, we focused on 14 out of 25 high-confident cell types that have more than 30 individuals after quality controls: ciliated epithelia cells, ENS glia, epicardial fat cells, hepatoblasts, intestinal epithelia cells, myeloid cells, neuroblasts, neurons, retinal neurons, retinal pigment cells, squamous epithelial cells, stromal cells, vascular endothelial cells, and iPSCs.

We included sex, the first two genotype principal components (PCs), and expression PCs as covariates. We computed expression PCs from the gene expression matrix for each cell type and condition to capture hidden technical or biological factors of expression variability. The number of PCs was selected and included in the model as covariates based on the elbow method and the Buja and Eyuboglu (BE) algorithm, which also approximately maximized cis-eGene discovery. We computed the correlation of the selected PCs and the other covariates and removed those that were highly correlated with any of the selected PCs (R^2^ > 0.9).

The mapping window was defined as 100kb up- and down-stream of the transcription start site. We output all results by setting the *pvOutputThreshold*=1. To identify genes with at least one significant eQTL (cis-eGenes), the top nominal p-value for each gene was selected and corrected for multiple testing at the gene level using eigenMT^63^. eigenMT is an efficient multiple hypothesis testing correction method that estimates the effective number of tests using a genotype correlation matrix and then applying Bonferroni correction. Genome-wide significance was then determined by computing Benjamini-Hochberg FDR on the top eigenMT-corrected p-value for each gene.

### Multivariate adaptive shrinkage

To combat the issue of incomplete power, we used mashr^64^ to estimate sharing between conditions, and to identify treatment-induced response effects in each cell type. By jointly analyzing genetic effects across conditions, mashr increases power and improves effect-size estimates, thereby allowing for greater confidence in effect sharing and estimates of condition-specificity. We ran mashr using the output of eQTL mapping from MatrixEQTL across 14 cell types in control and each treatment group in pairs (28 effects for each SNP). We included a total of 9,717 genes that are tested in each cell type and condition combination. No effect size is *post hoc* set to 0 for mashr. To run mashr, we first estimated residual correlation structure on all independent gene-SNP pairs (around 100,000) from an intersection of all tested SNPs and pruned reference SNPs (--indep-pairwise 50 5 0.2). We obtained a similar correlation structure when we randomly selected 100,000 gene-SNP pairs regardless of linkage disequilibrium. Next, to estimate data-driven covariance matrices, we selected the top SNP with the smallest p-value across all 28 conditions for each gene as a strong set of 9717 gene-SNP pairs. For genes that have multiple selected SNPs with the same smallest p-value, we randomly selected one SNP. Using this strong set, we obtained empirical covariance matrix, PCA-derived matrix, and matrices computed by empirical Bayes matrix factorization via flashr using its default settings.

Next, we learned the underlying parameters on the list of data-driven covariance matrices using extreme deconvolution. We fit the mash model to our random set of independent gene-SNP pairs (that were used for estimating residual correlations) using both data-driven and canonical covariances. Finally, we computed posterior statistics for the strong set of 9717 gene-SNP pairs.

For each cell type, gene-SNPs and LFSR < 0.1 in control or treatment were designated as significant eQTLs. To investigate sharing of the eQTLs between control and treatment, we assessed sharing of effects by magnitude (effects have similar magnitude within a factor of 1.5) for the SNPs that are significant eQTLs within each cell type separately. Response eQTLs are SNPs that are significant eQTLs in control or treatment for a given cell type and the fold change of effects is more than a factor of 1.5. A response eGene is a gene with at least one response eQTL. Non-response eQTLs are SNPs that are significant eQTLs in at least one condition for a given cell type and the fold change of effects is less than a factor of 1.5. A non-response eGene is a gene with at least one eQTL but no response eQTLs.

### Dynamic eQTL mapping

First, we aggregated cells into pseudobulk samples by grouping cells into 15 pseudotime bins of equal range in each trajectory of each condition. We removed donors with fewer than 5 cells for a given trajectory. Then, we merged the bin with fewer than 100 or total number of cells divided by 15 (whichever is smaller) into the next bin. We iteratively merge the bins until no bin has too few cells. If the last bin has too few cells, merge it into the second to the last bin. We performed similar merging procedure by individual to make sure each bin has at least 20 individuals. Then, we normalized pseudobulk expression by individual level in each bin and used the filtered genotypes as processed in the *molecular cis-eQTL mapping* section.

Following prior approaches^4,19,65^, we run PCA on cell lines to capture latent factors that account for broad expression differences across individuals during differentiation. PCA was conducted using the *NIPALS algorithm*^66^, which accommodates missing data. In our regression model, we included sex and the first 10 cell line PCs, along with their interactions with pseudotime, as covariates to adjust for lineage-specific effects. Additionally, we included the top 2 genotype PCs as covariates, without their interactions with pseudotime, to control for population structure.

We implemented dynamic eQTL mapping with TensorQTL^33^ using the following linear model:

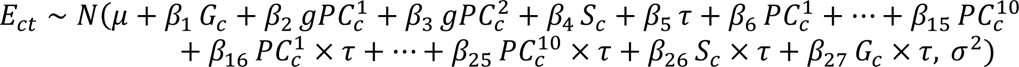

Where *c* indexes donors and *t* indexes pseudotime bins, *E_ct_* represents the normalized expression of the sample, *G* represents genotype, *gPC* represents genotype PCs, *PC_k_* represents kth cell line PCs, Sc represents the sex of donor c, and τ represents median pseudotime value of cells in each pseudotime bin.

We assessed dynamic eQTLs by modeling the interaction between genotype and pseudotime (*β*_27_). To correct for the multiple variant-level tests performed per gene, we applied eigenMT^63^, followed by the Benjamini-Hochberg procedure on the minimum eigenMT-adjusted p-value per gene to control the genome-wide FDR at 0.1.

To define significant dynamic eQTLs, we first identified the gene whose eigenMT-adjusted p-value was closest to the genome-wide FDR threshold and used that p-value as the significance cutoff. All variant-gene pairs with adjusted p-values below this threshold were retained as significant dynamic eQTLs.

For replication analysis, we evaluated the consistency of our neural dynamic eQTLs using summary statistics from J. Popp and K. Rhodes et al^19^. We estimated the *π₁* replication rate using the *qvalue* package, focusing on the neural dynamic eQTL identified in our control samples.

We categorized the significant dynamic eQTLs into early, late, or switch classes, following established criteria^4,19,65^. An eQTL was labeled early if its effect size decreased across pseudotime, late if the effect size increased, and switch if the direction of the effect reversed. To make these classifications, we used predicted effect sizes at the trajectory endpoints from the fitted linear model produced by TensorQTL. If the direction remained consistent, the classification was based on whether the magnitude decreased (early) or increased (late). If the effect size changed sign and both endpoint magnitudes exceeded a threshold of 1, the variant was considered a switch-effect eQTL.

### Enrichment in regulatory elements

We annotate the variants using signatures of cis-regulatory elements (cCREs) from ENCODE^36^. In the ENCODE project, epigenetic signatures and proximity to TSSs were integrated to categorize cCREs. We used all types of cCREs in our analysis but focused on promoter domains and enhancer domains in the results. Promoter-like signatures (PLS) must meet the following two criteria: 1) fall within 200 bp (center to center) of an annotated GENCODE TSS or experimentally derived TSS and 2) have high chromatin accessibility and H3K4me3 signals.

Enhancer-like signatures (ELS) have high chromatin accessibility and H3K27ac signals. If they are within 200 bp of a TSS they must also have low H3K4me3 signal. The subset of the enhancers within 2 kb of a TSS are denoted as TSS proximal (pELS), while the remaining subset is denoted TSS distal (dELS). We obtained all human cCREs (GRCh38/hg38) across all cell types from ENCODE’s Registry beta version. We annotated all SNPs tested in the eQTL mapping if it is located within an cCRE. Enrichment tests were performed in each cell type and treatment using Fisher’s exact test.

### Comparison of gene-level features

We compared genic features of response eGenes, non-response eGenes, and all eGenes in each cell type. We calculated the TSS count of a given gene by the number of promoter peaks measured by the FANTOM5 project using Cap Analysis of Gene Expression (CAGE)^38^. We estimated enhancer activity of a given gene by computing the accumulative enhancer length across 131 cell or tissue types predicted based on the activity-by-contact (ABC) model (ABC score > 0.05)^42^. We defined loss-of-function (LoF)-intolerant genes by pLI score > 0.9 using the statistic from gnomAD.v4.1.0^67^. For Gene Ontology (GO) analysis, we focused on 41 broadly unrelated GO terms that are used in Mostafavi et al^35^. For enrichment in transcription factors (TFs), we used the official list of human TFs (v1.01) from Lambert et al. which includes 1639 human TFs^68^. For all the analyses, we only included the genes that were shared between the testing set for eQTL mapping in this study (n=9717) and the reference set of the interest. Enrichment tests were performed using Fisher’s exact test.

### Colocalization

We compiled GWAS summary statistics from Sakaue et al.^44^ and Fair et al.^45^ The first study contains a GWAS atlas for 220 human phenotypes from BioBank Japan, UK Biobank, and FinnGen (n_total=807,000). The second study compiled summary statistics for 45 blood and immune-related traits from GWAS Catalog. We used HyPrColoc to determine whether eQTL and trait associations are explained by the same causal variant. HyPrColoc is an efficient deterministic Bayesian divisive clustering algorithm that uses GWAS summary statistics to detect colocalization across vast numbers of traits simultaneously^46^. We used the default setting of prior probability = 1×10^-4^ and a conditional colocalization prior = 0.025. We defined a significant colocalization as having a posterior probability larger than 0.5.

---

**Figure S1.**
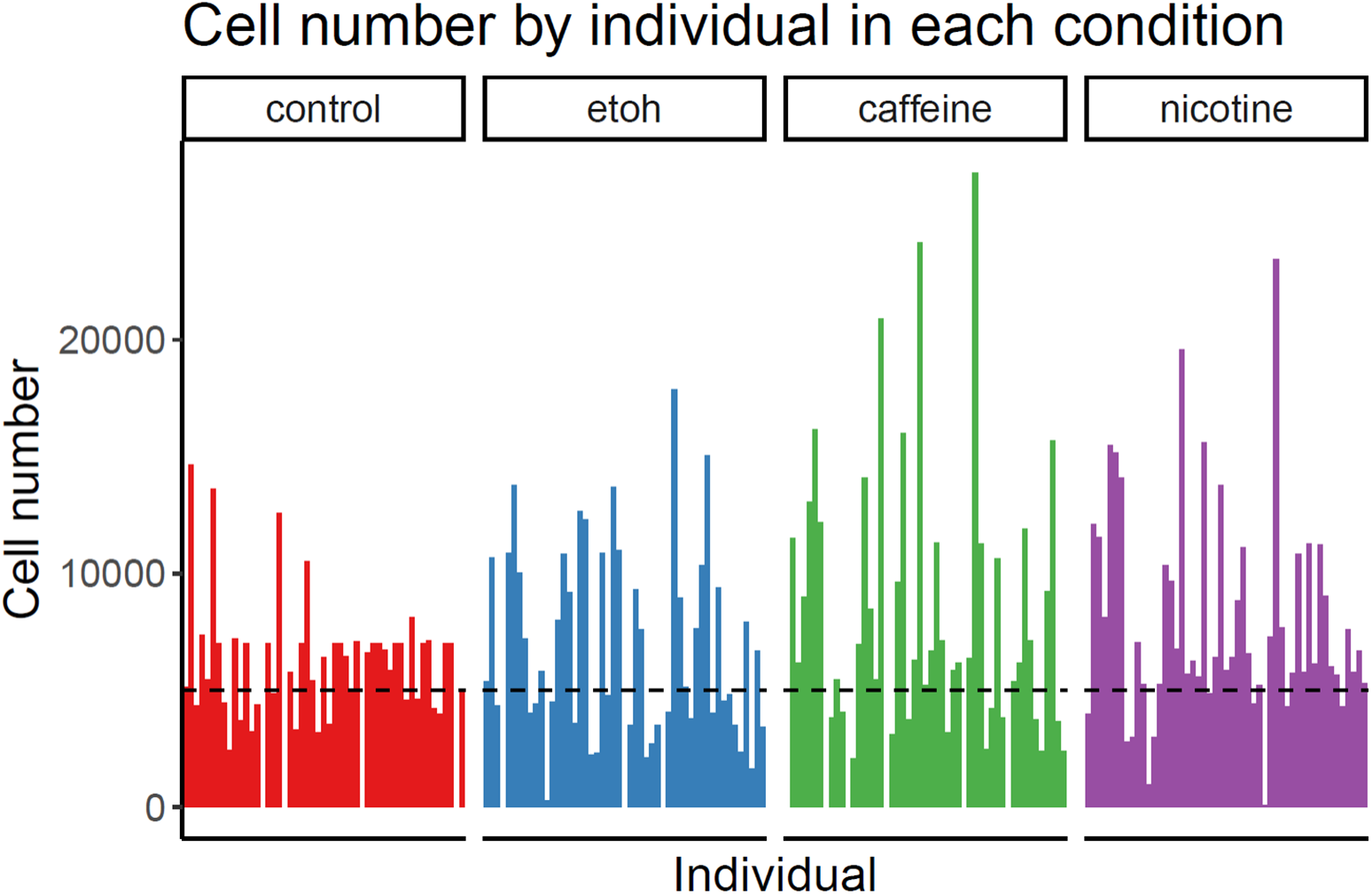
Summary of cell number in each individual line and condition.

**Figure S2.**
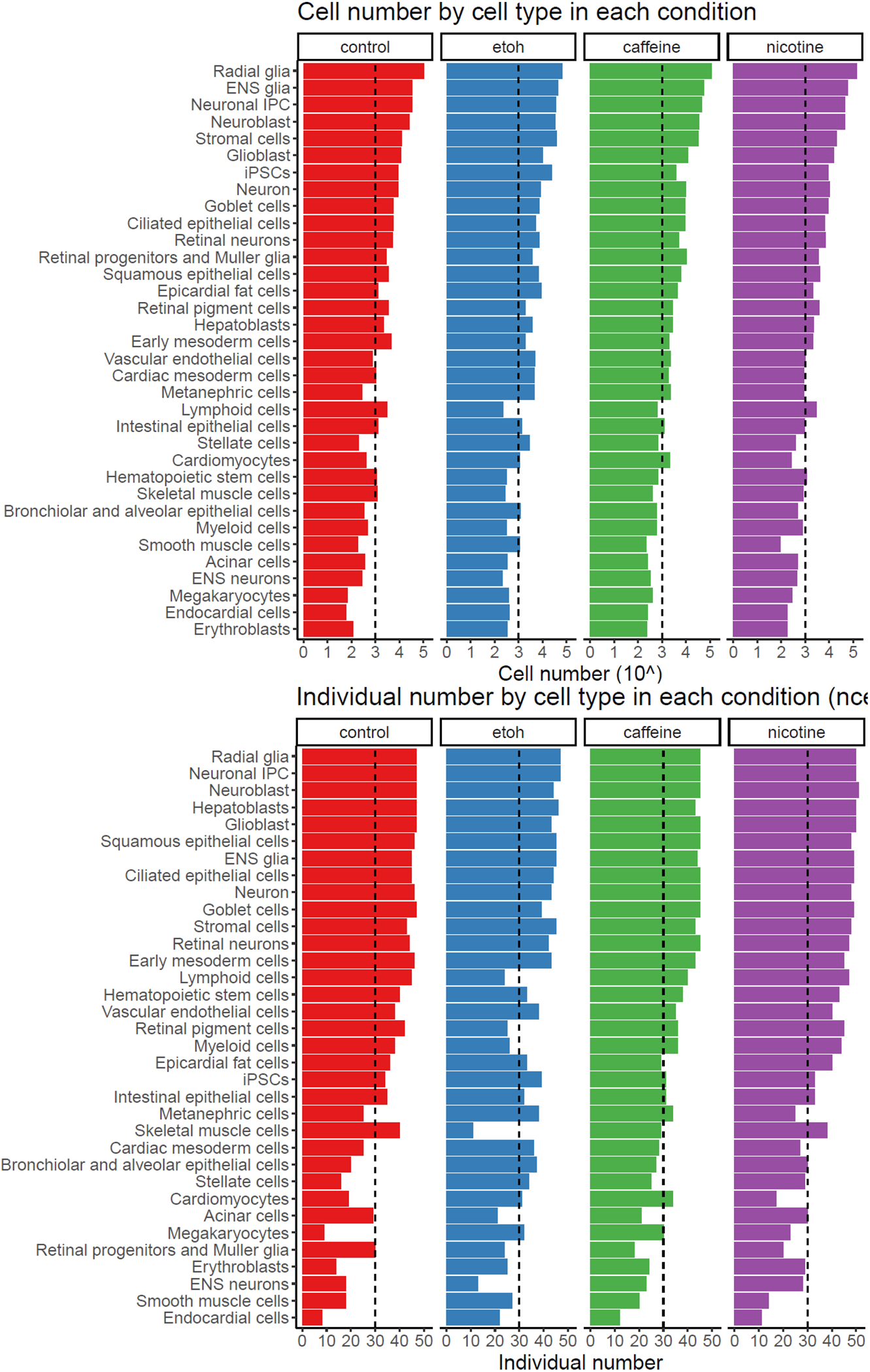
Summary of cell number and individual number in each cell type and condition.

**Figure S3.**
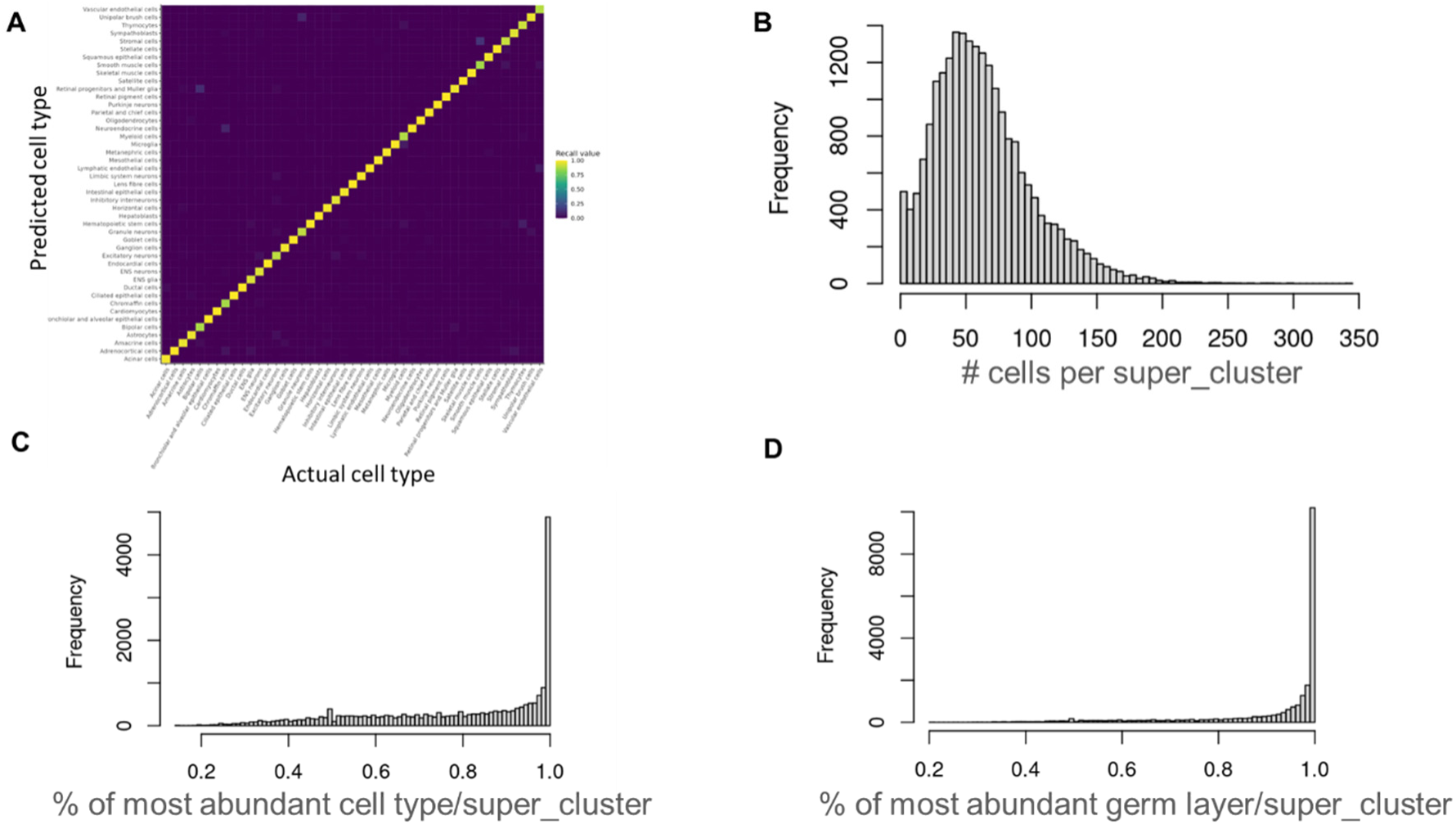
Validation of cell type annotation approach and results. (A) Confusion matrix for intradataset cell type cross-validation. We split the fetal cell atlas into training and test sets where we used the training set to generate a reference marker list and the test set for cell type prediction. (B) Distribution of cell number across 22K super clusters generated by unsupervised clustering with a high resolution, with an average of 60 cells per cluster across 1.4M cells in our study. (C-D) Histogram of the proportion of most abundant assigned cell type (C) or germ layer (D) in each super cluster. Cells in the majority of super clusters have a homogenous (>80%) cell type/germ layer assignment.

**Figure S4.**
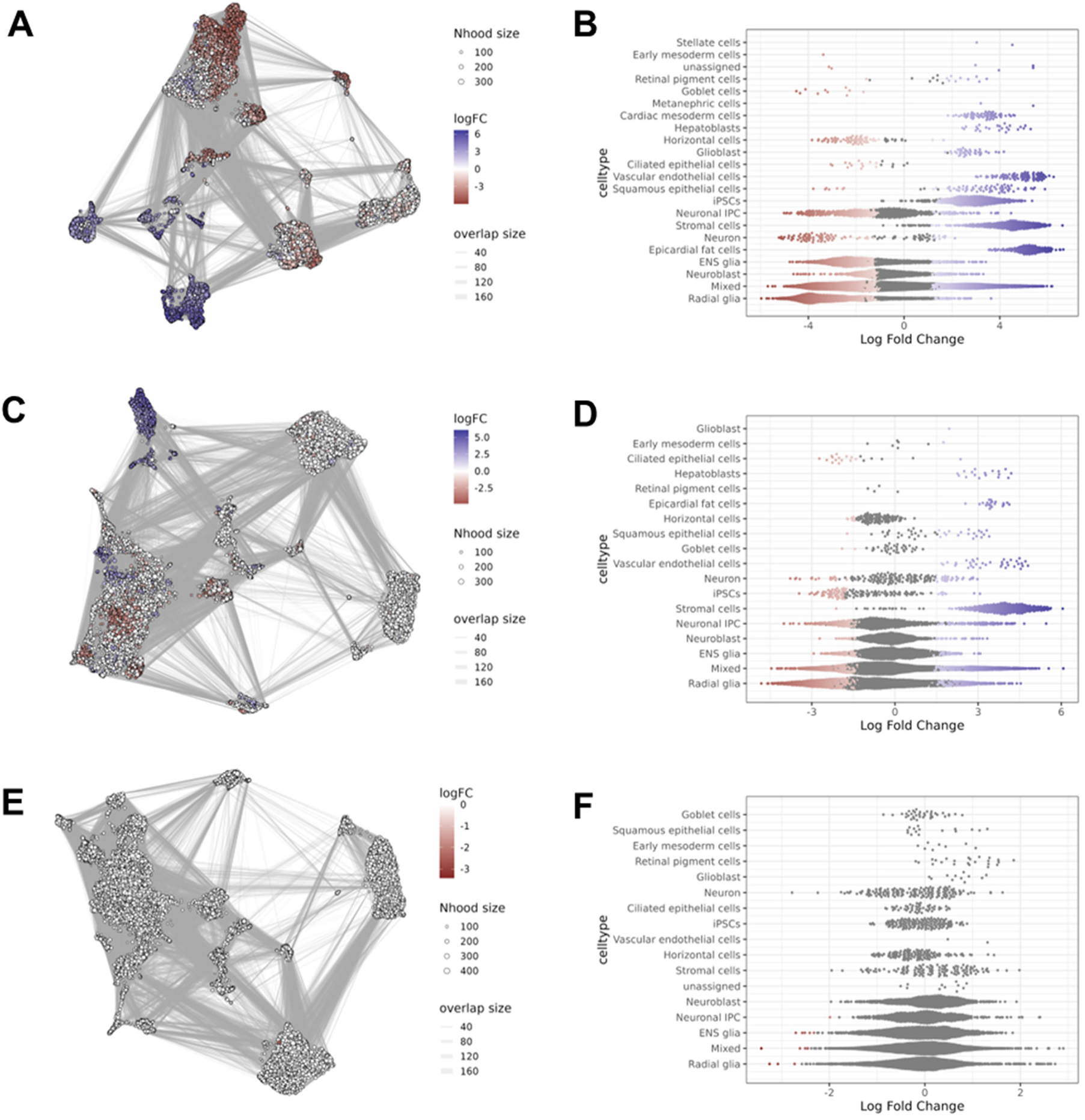
Differential abundance in response to treatment. (A, C, E) Visualization of differential abundance on a KNN graph. Nodes are neighborhoods, colored by their log fold change between control and ethanol (A), caffeine (C), and nicotine (E), respectively. Non-differential abundance neighborhoods (FDR>5%) are colored white, and sizes correspond to the number of cells in each neighborhood. Graph edges depict the number of cells shared between neighborhoods (B, D, F) Distribution of log fold change between treatment and control in neighborhoods containing cells from different cell type. Differential abundance neighborhoods at FDR<5% are colored. A neighborhood is assigned to a cell type if >75% cells within the neighborhood from the same predefined cell type.

**Figure S5.**
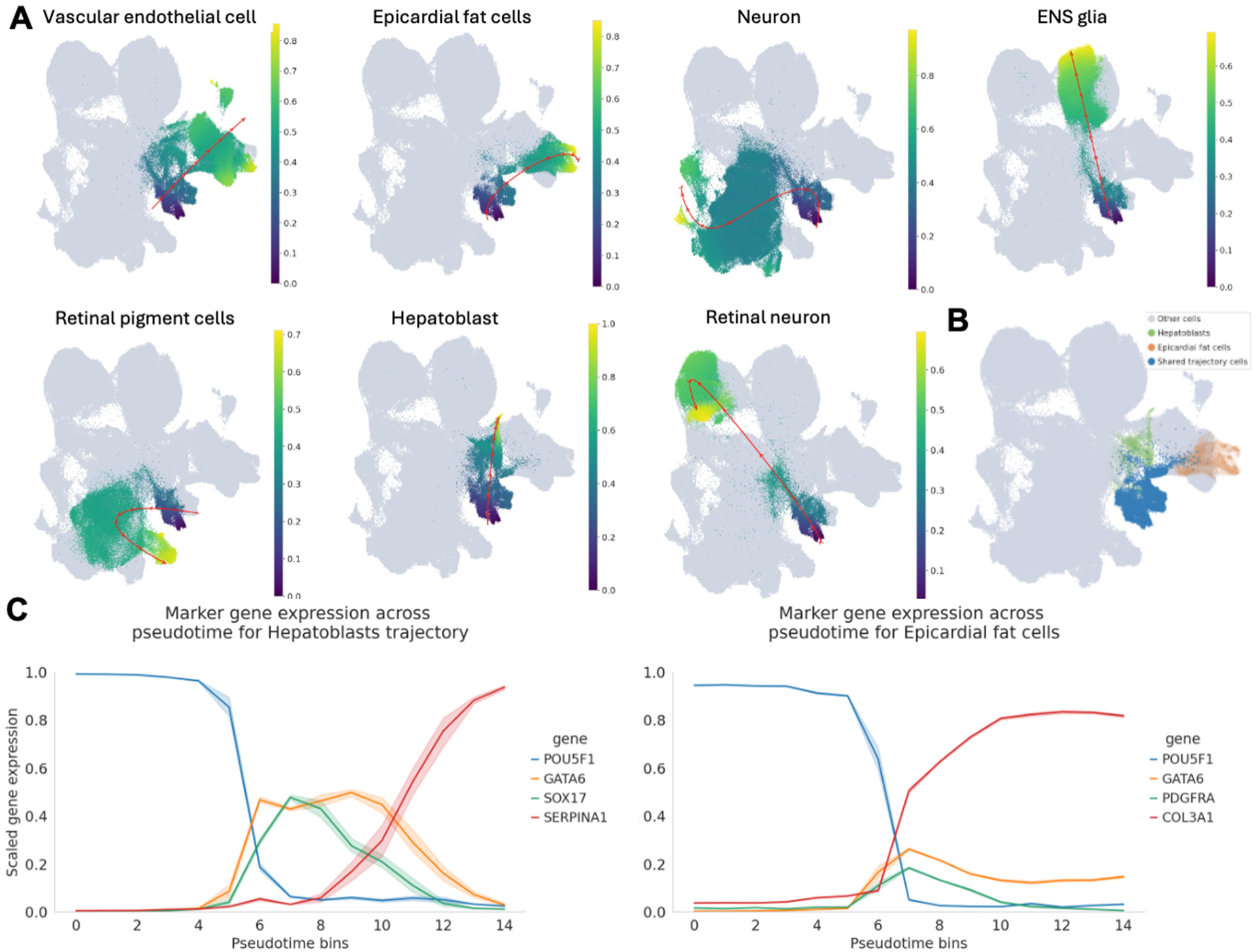
Full trajectories and trajectory validation. **(A)** Pseudotime distributions for all inferred trajectories. Color bars represent pseudotimes of cells assigned to each trajectory. Cells not assigned to the trajectory are shown in gray. **(B)** Hepatoblast and epicardial fat cell trajectories share intermediate mesendoderm cells. Cells assigned in hepatoblast only are shown in green, cells assigned in epicardial fat cells only are orange, and cells shared by the two trajectories are shown in blue. Cells not assigned to either trajectory are shown in gray. **(C)** Marker gene expression for hepatoblast (left) and epicardial fat cell (right) trajectories. *POU5F1* is pluripotent cell marker gene, *GATA6* is the intermediate mesendoderm marker gene, *SOX17* is intermediate mesoderm marker gene, *PDGFRA* is intermediate endoderm marker gene, *SERPINA1* is hepatoblasts marker gene, and *COL3A1* is epicardial fat cell marker gene.

**Figure S6.**
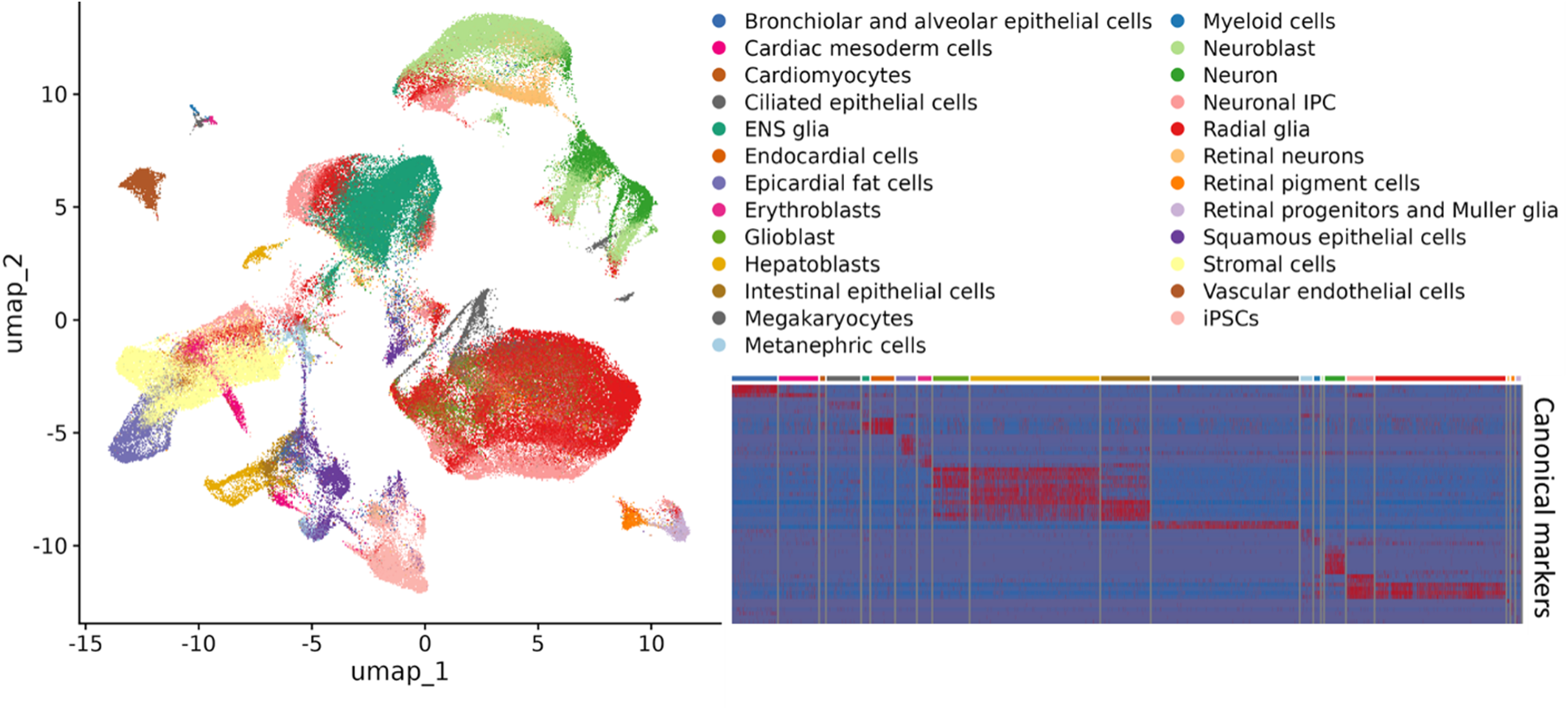
Validation of 25 high-confidence cell types. UMAP visualization of cell types in different colors and heatmap of expression of canonical markers for each cell type.

**Figure S7.**
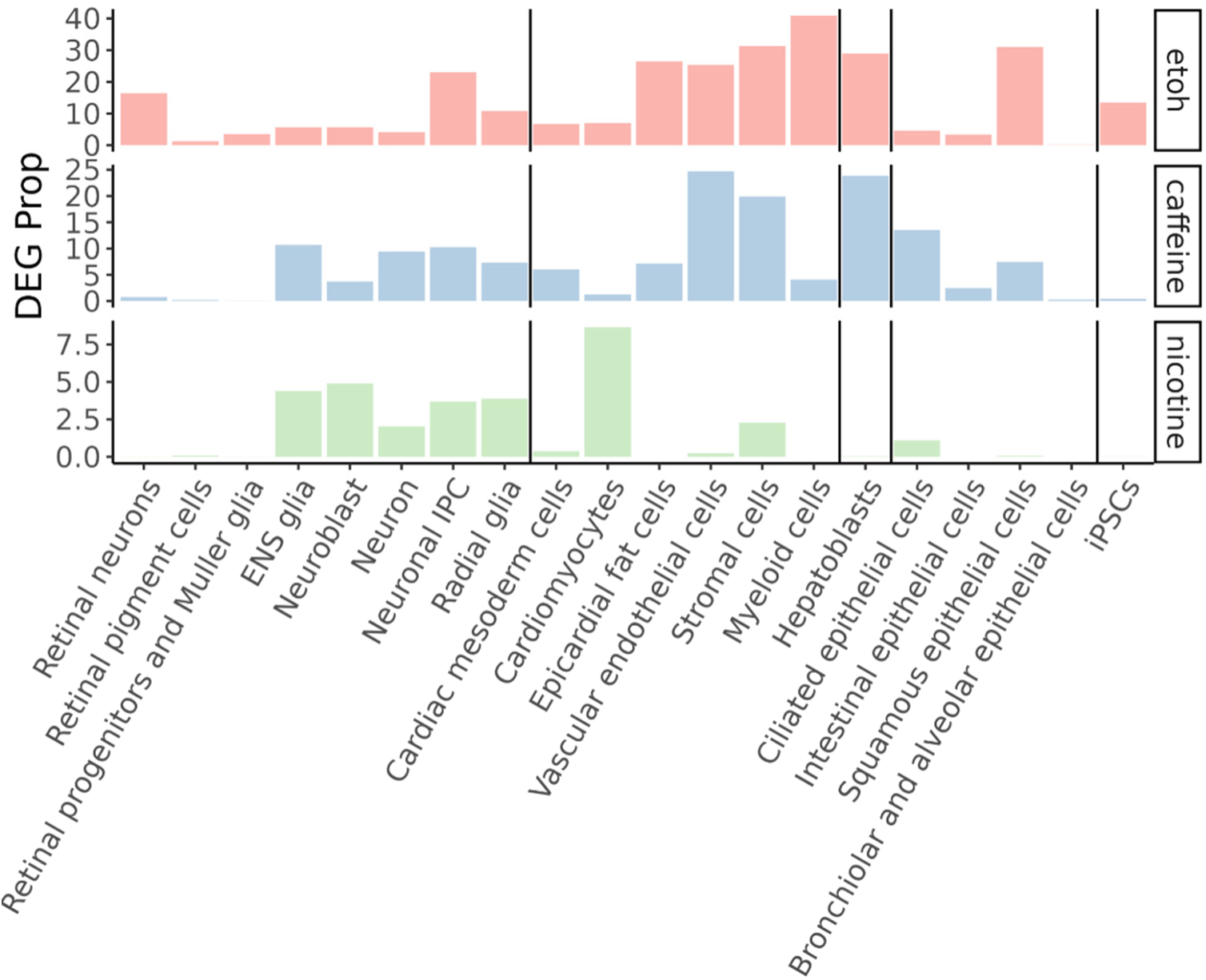
Proportion of treatment-induced differentially expressed genes in each cell type.

**Figure S8.**
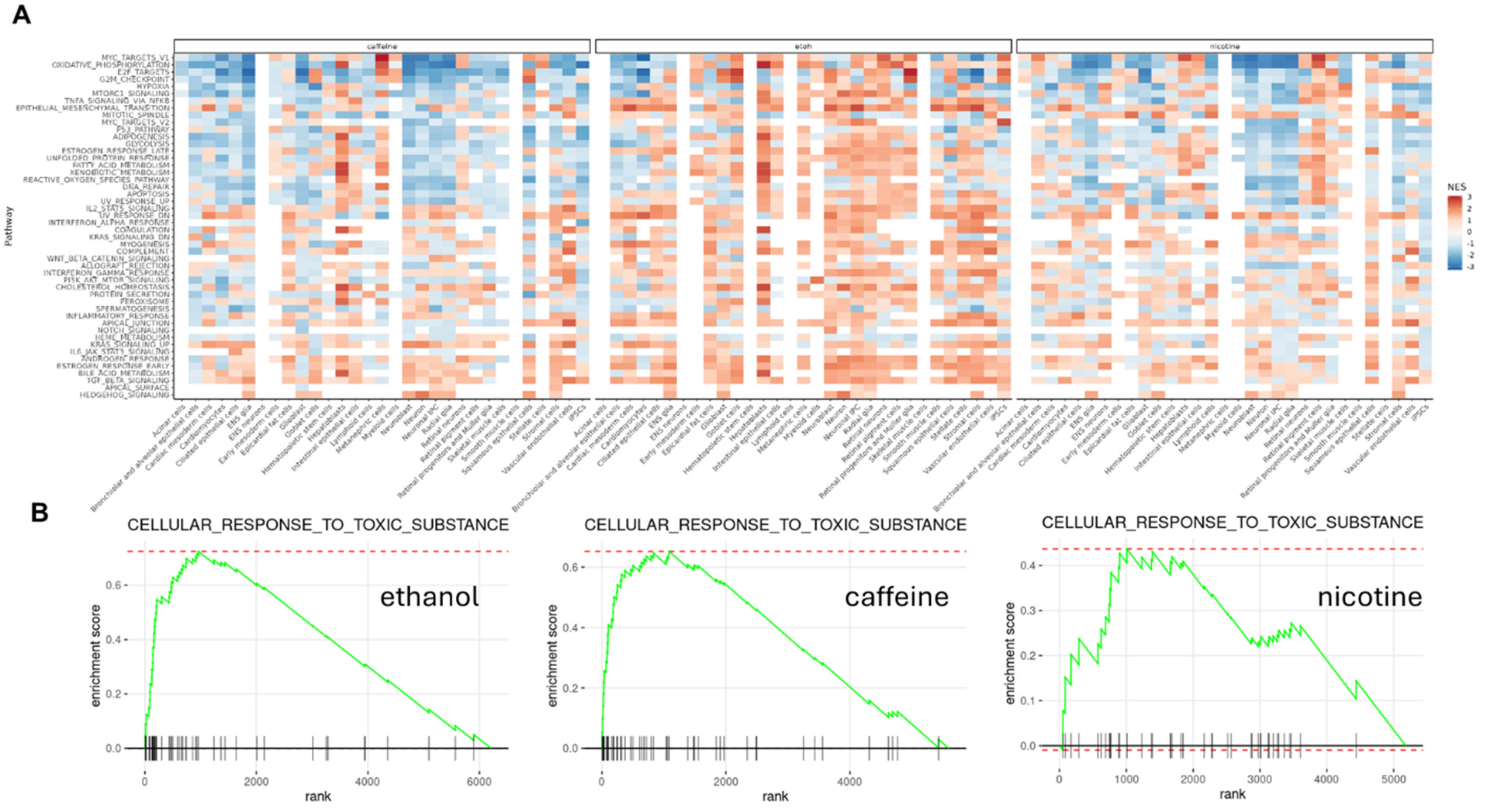
Pathway enrichment of differentially expressed genes in response to each treatment. (A) Heatmap of enrichment in 50 hallmark gene set. Normalized enrichment score (NES) indicates direction and strength of enrichment, with enrichment in red and depletion in blue. (B) Ranks of genes involved in cellular response to toxic substances by t-statistics of differential expression test in hepatoblasts in response to ethanol, caffeine, and nicotine, respectively.

**Figure S9.**
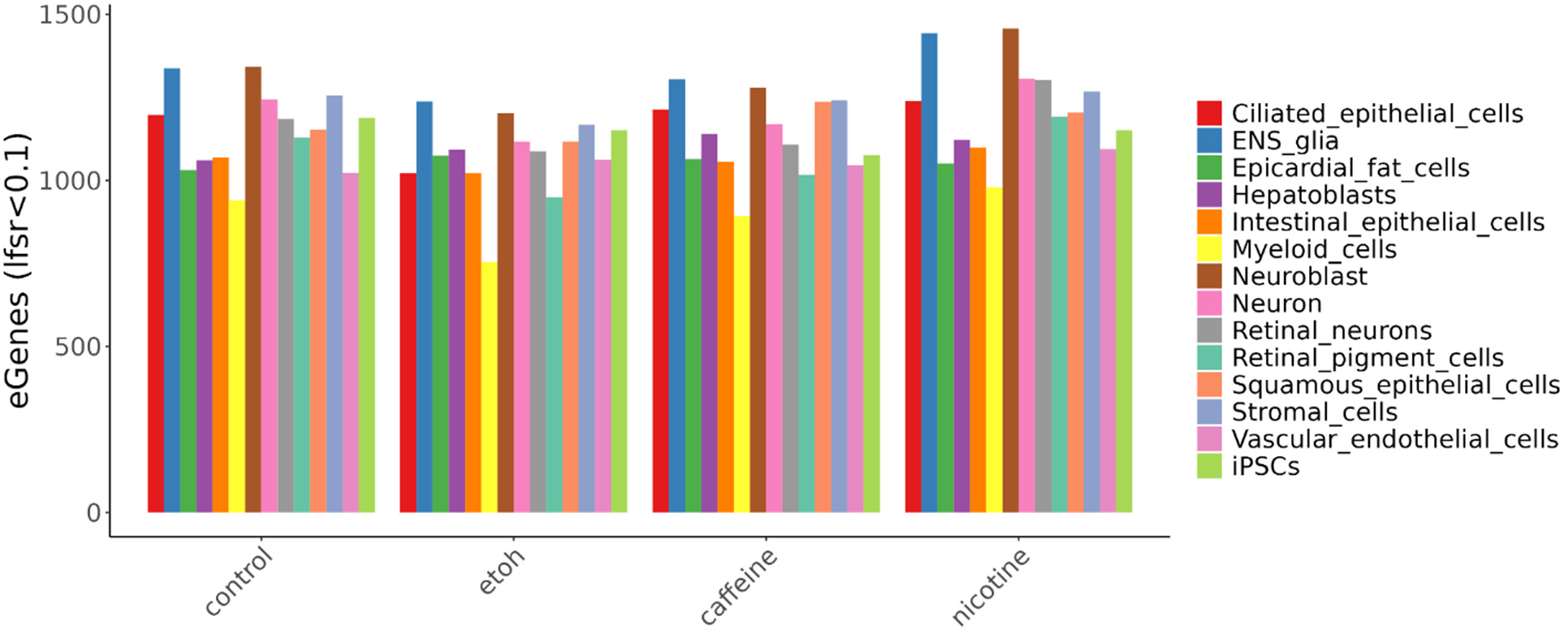
eGenes discovery in each cell type and condition.

**Figure S10.**
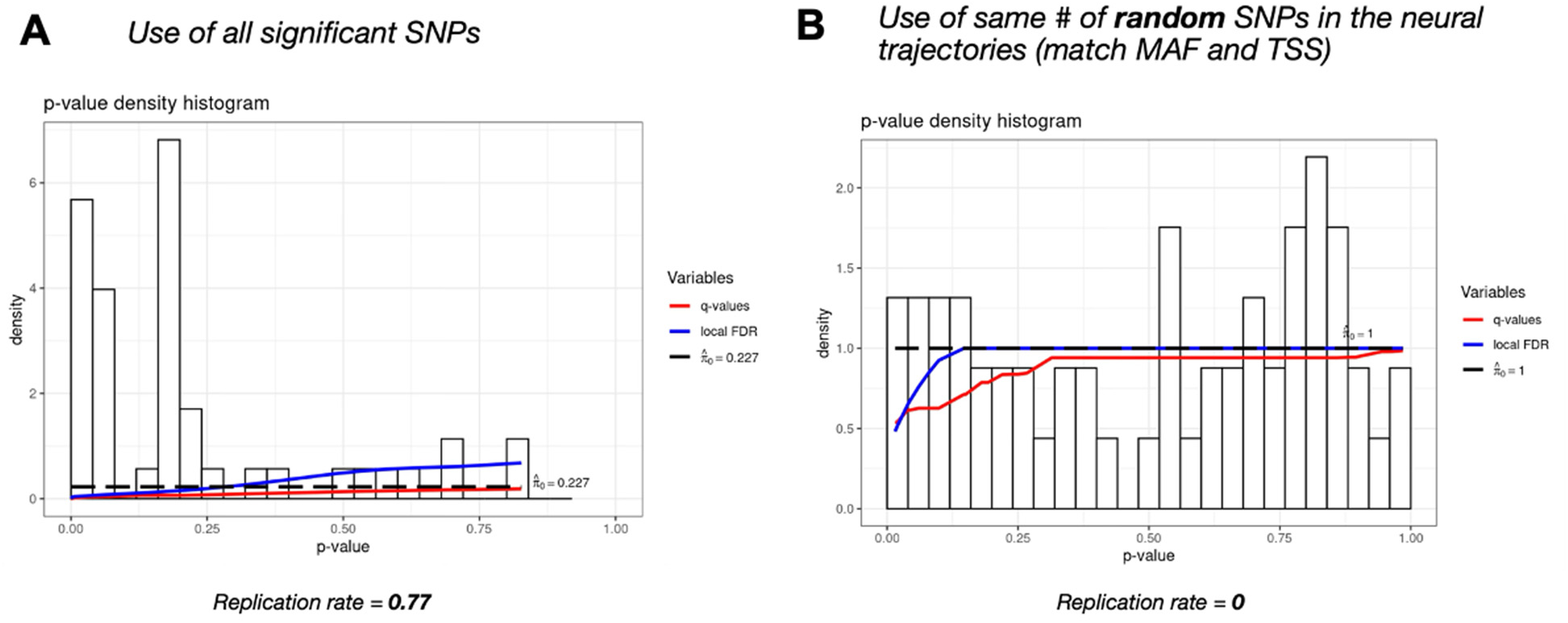
Replication of dynamic eQTLs. (A) Replication of control neural dynamic eQTLs in J. Popp and K. Rhodes et al. and (B) their null set of eQTLs with matched minor allele frequence (MAF) and transcription start site (TSS).

**Figure S11.**
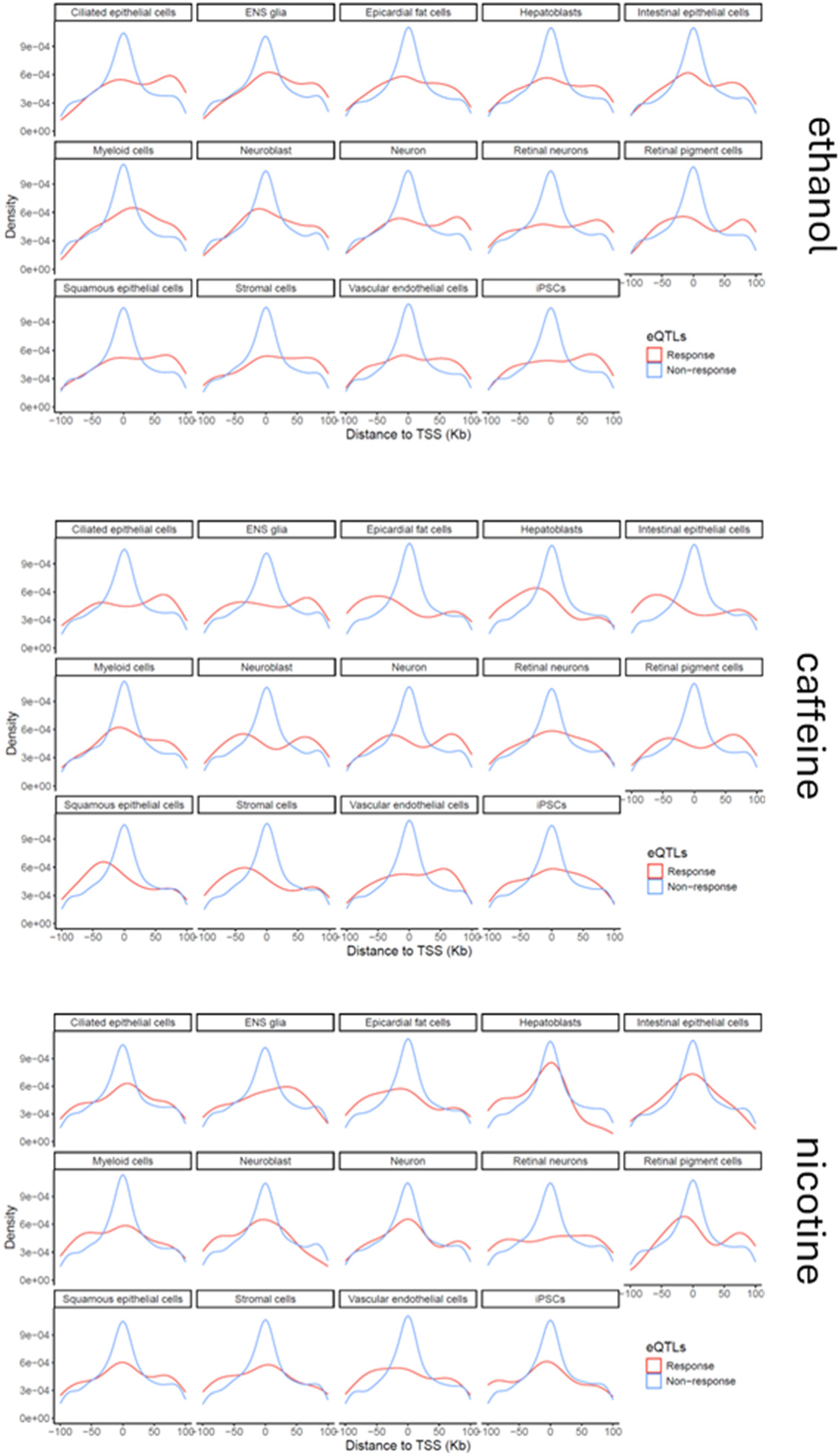
Distribution of distance to TSS for response eQTLs and non-response eQTLs in each cell type and treatment.

**Figure S12.**
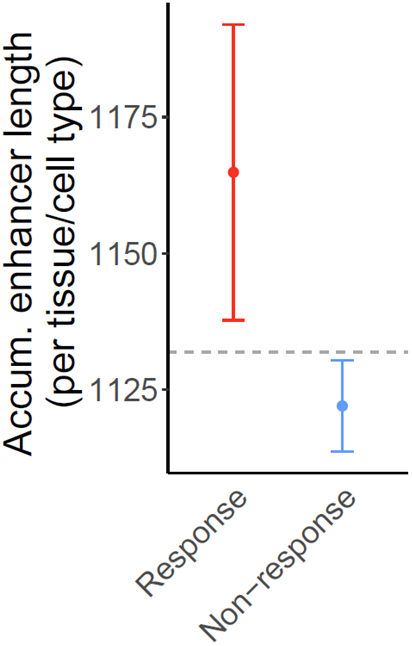
Average cumulative enhancer length per tissue/cell type.

**Figure S13.**
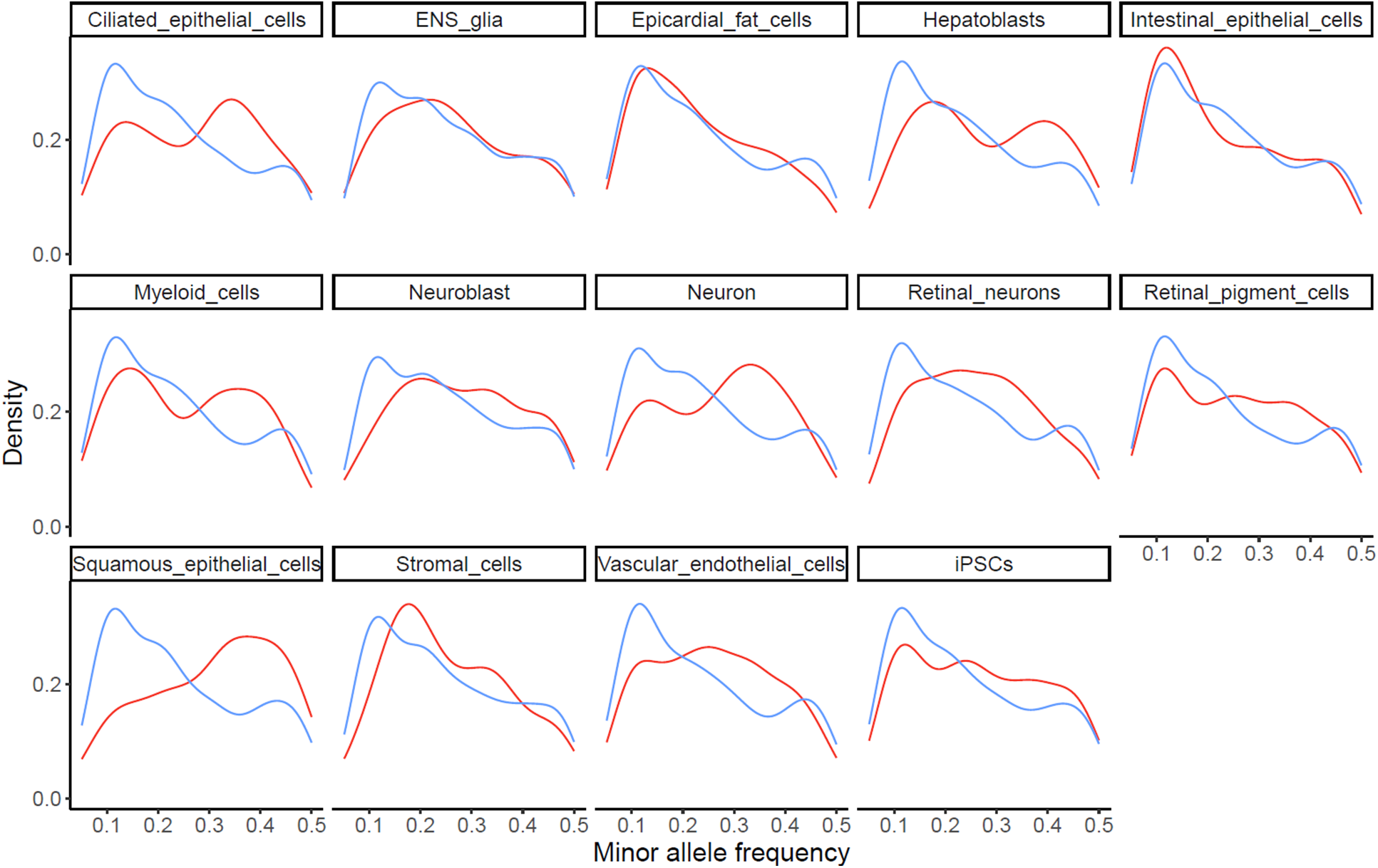
Distribution of minor allele frequency (MAF).

**Figure S14.**
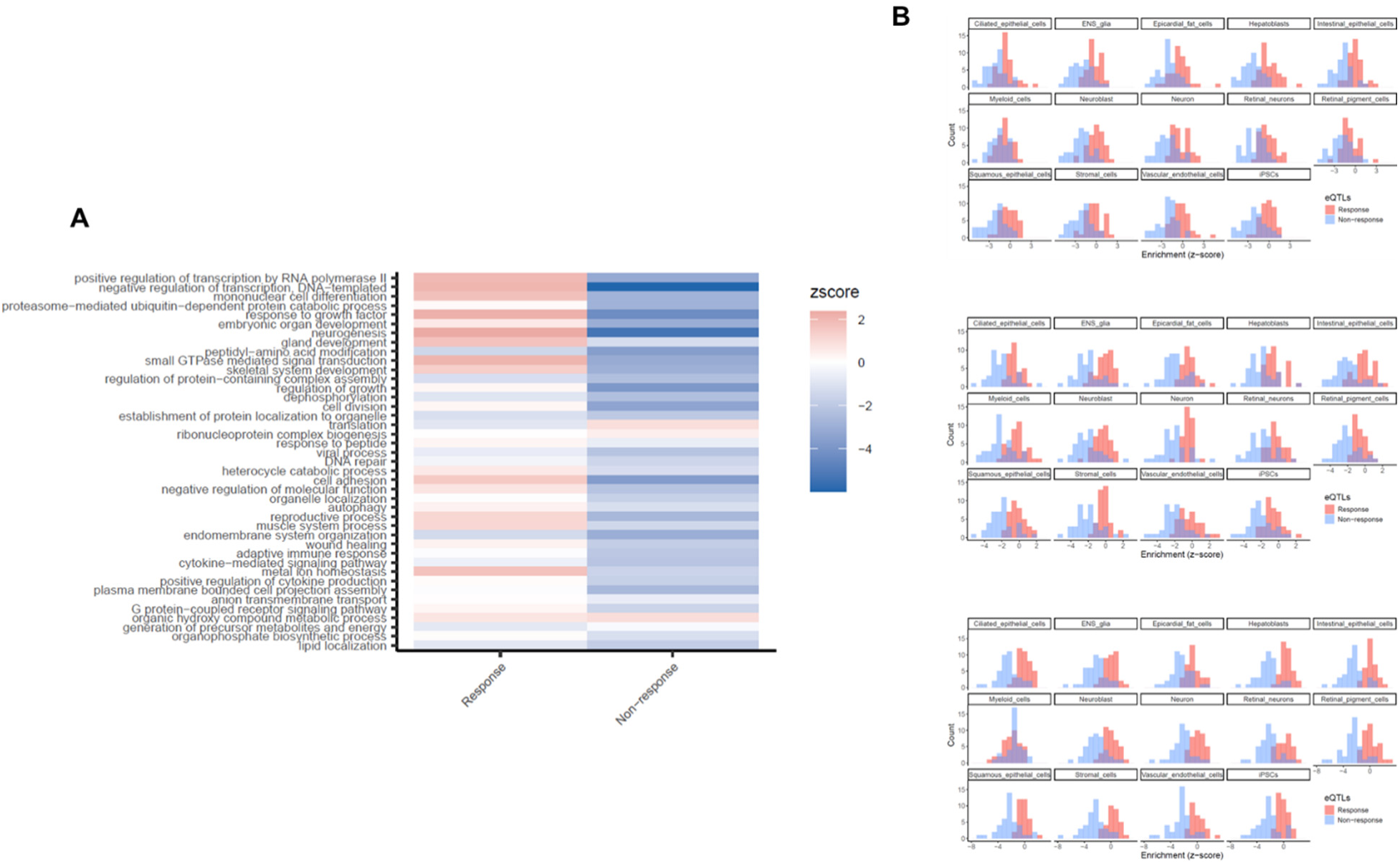
Enrichment of response eGenes and non-response eGenes in biological processes. (A) Enrichment of response eGenes and non-response eGenes among genes in 41 broadly defined Gene Ontology (GO) categories. The GO categories (y-axis) are sorted based on the average pLI value of the corresponding genes within each category. The color map represents enrichment (red) or depletion (blue) by enrichment Z scores. (B) Distribution of enrichment Z scores across 41 GO terms for response eGenes (red) and non-response eGenes (blue). From top to bottom: ethanol, caffeine, and nicotine.

**Figure S15.**
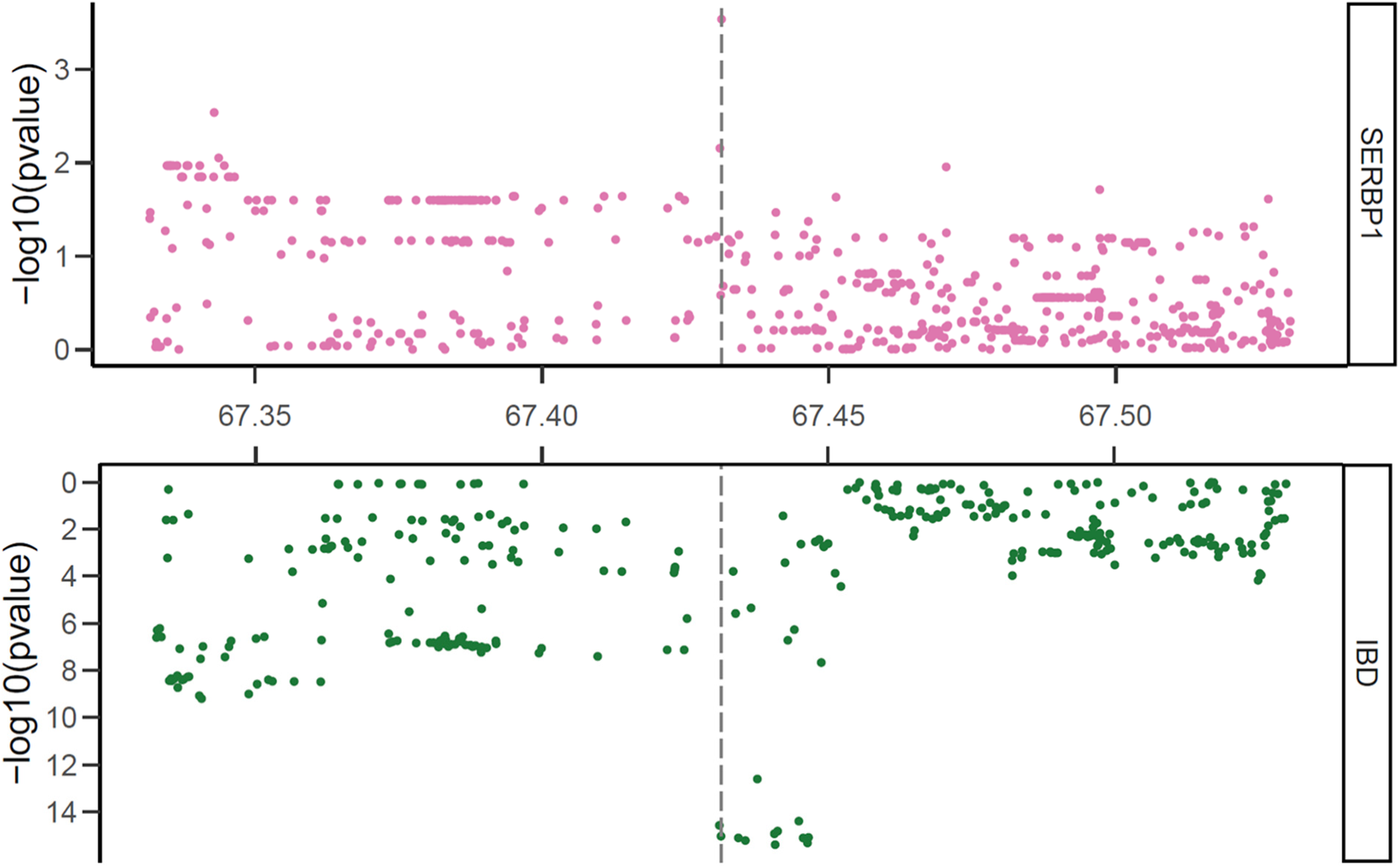
Colocalization of a response eQTL with IBD. Visualization of an example of a caffeine-response eQTL (chr1:67431276:G_T) for *SERBP* in intestinal epithelial cells colocalized with a variant associated with IBD. The top panel (pink) shows significance levels of variants evaluated as eQTLs for the given gene expression including all variants within 100kb of the transcription start site, and the bottom panel (green) shows significance levels of variants tested for association with the trait(s) within the same region. Vertical lines depict the genomic location of the candidate colocalized variant.

**Figure S16.**
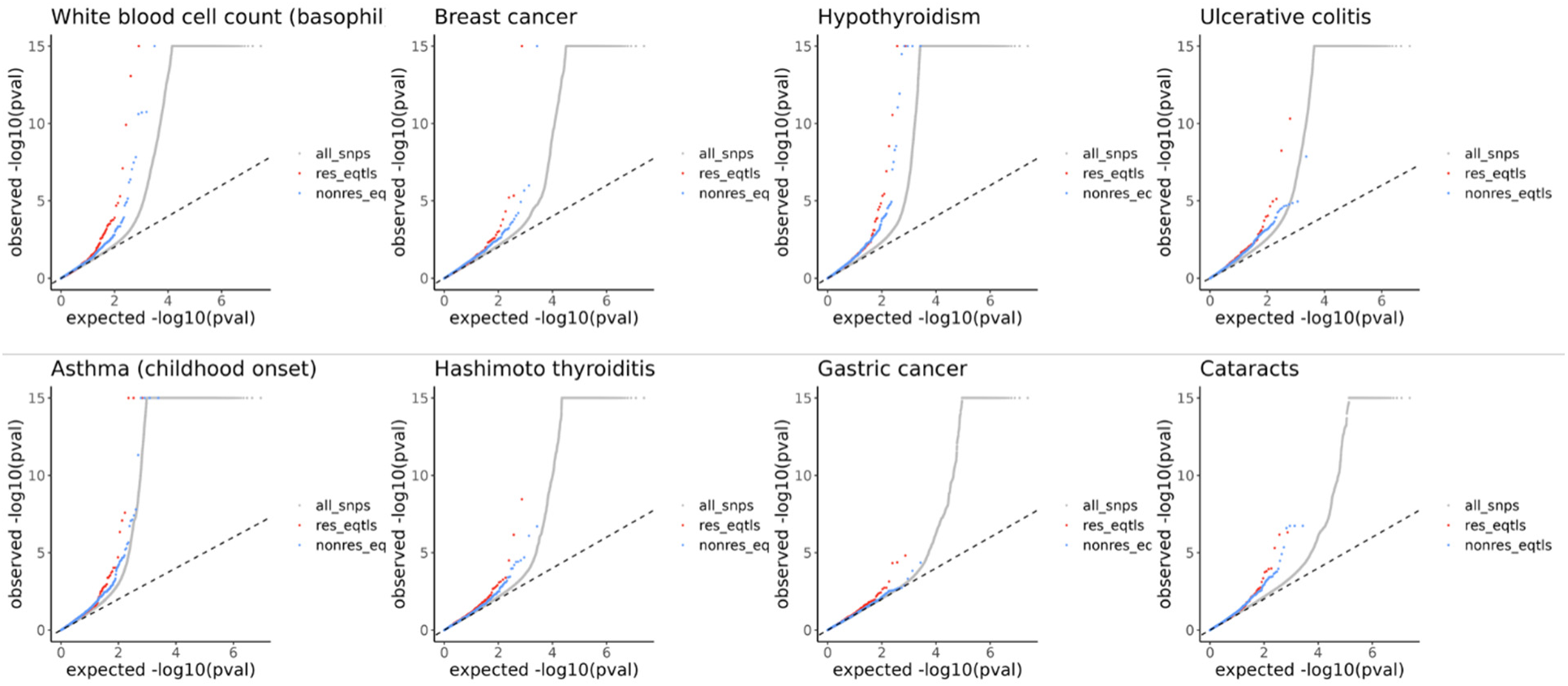
Examples of stronger enrichment of GWAS P-values in response eQTLs.

**Figure S17.**
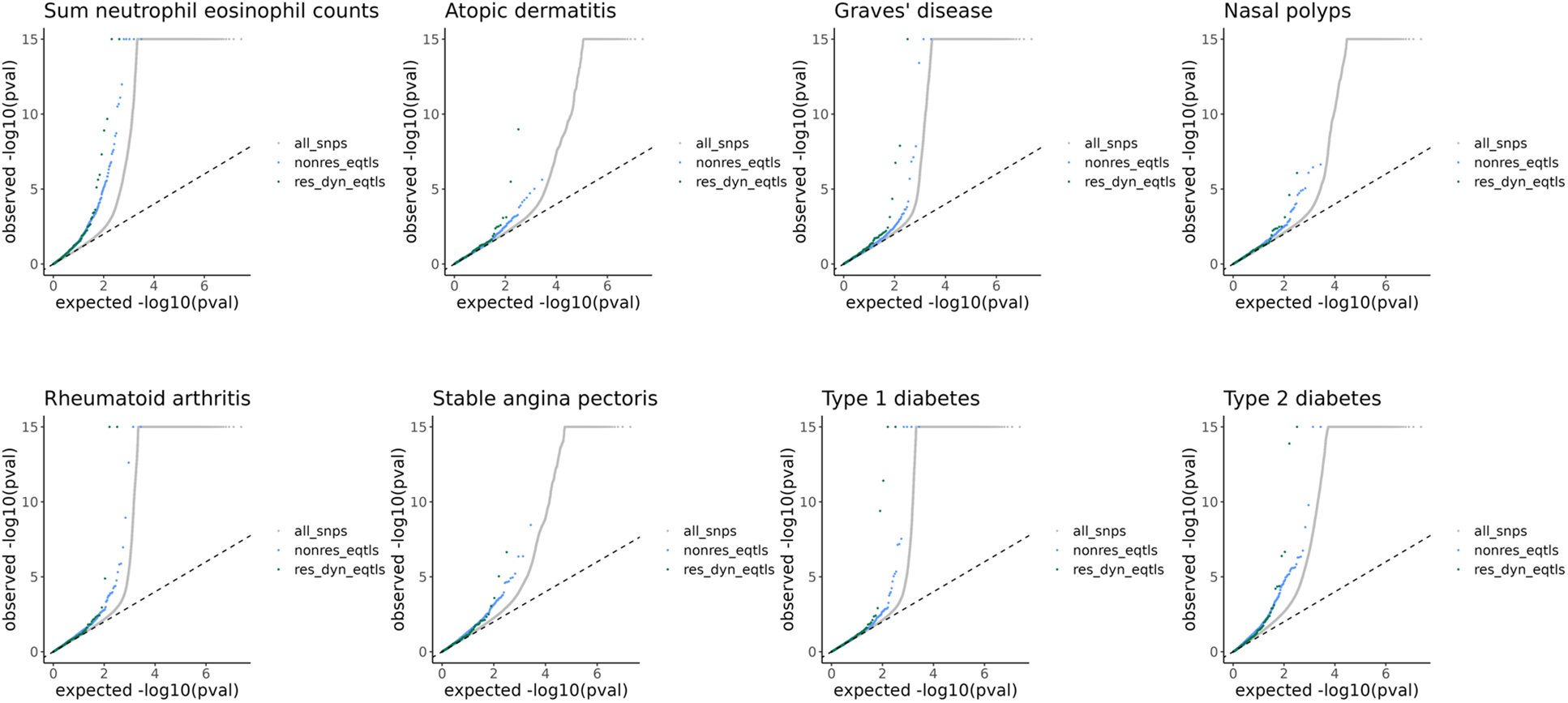
Examples of stronger enrichment of GWAS P-values in dynamic response eQTLs.

## Reference

1. Albert, F.W., and Kruglyak, L. (2015). The role of regulatory variation in complex traits and disease. Nat. Rev. Genet. 16, 197–212. 10.1038/nrg3891.

2. THE GTEX CONSORTIUM (2020). The GTEx Consortium atlas of genetic regulatory effects across human tissues. Science 369, 1318–1330. 10.1126/science.aaz1776.

3. Nathan, A., Asgari, S., Ishigaki, K., Valencia, C., Amariuta, T., Luo, Y., Beynor, J.I., Baglaenko, Y., Suliman, S., Price, A.L., et al. (2022). Single-cell eQTL models reveal dynamic T cell state dependence of disease loci. Nature 606, 120–128. 10.1038/s41586-022-04713-1.

4. Strober, B.J., Elorbany, R., Rhodes, K., Krishnan, N., Tayeb, K., Battle, A., and Gilad, Y. (2019). Dynamic genetic regulation of gene expression during cellular differentiation. Science. 10.1126/science.aaw0040.

5. Ward, M.C., Banovich, N.E., Sarkar, A., Stephens, M., and Gilad, Y. (2021). Dynamic effects of genetic variation on gene expression revealed following hypoxic stress in cardiomyocytes. eLife 10, e57345. 10.7554/eLife.57345.

6. Fairfax, B.P., Humburg, P., Makino, S., Naranbhai, V., Wong, D., Lau, E., Jostins, L., Plant, K., Andrews, R., McGee, C., et al. (2014). Innate immune activity conditions the effect of regulatory variants upon monocyte gene expression. Science 343, 1246949. 10.1126/science.1246949.

7. Gutierrez-Arcelus, M., Baglaenko, Y., Arora, J., Hannes, S., Luo, Y., Amariuta, T., Teslovich, N., Rao, D.A., Ermann, J., Jonsson, A.H., et al. (2020). Allele-specific expression changes dynamically during T cell activation in HLA and other autoimmune loci. Nat. Genet. 52, 247–253. 10.1038/s41588-020-0579-4.

8. Ota, M., Nagafuchi, Y., Hatano, H., Ishigaki, K., Terao, C., Takeshima, Y., Yanaoka, H., Kobayashi, S., Okubo, M., Shirai, H., et al. (2021). Dynamic landscape of immune cell-specific gene regulation in immune-mediated diseases. Cell 184, 3006–3021.e17. 10.1016/j.cell.2021.03.056.

9. Schmiedel, B.J., Gonzalez-Colin, C., Fajardo, V., Rocha, J., Madrigal, A., Ramírez-Suástegui, C., Bhattacharyya, S., Simon, H., Greenbaum, J.A., Peters, B., et al. (2022). Single-cell eQTL analysis of activated T cell subsets reveals activation and cell type–dependent effects of disease-risk variants. Sci. Immunol. 7, eabm2508. 10.1126/sciimmunol.abm2508.

10. Soskic, B., Cano-Gamez, E., Smyth, D.J., Ambridge, K., Ke, Z., Matte, J.C., Bossini-Castillo, L., Kaplanis, J., Ramirez-Navarro, L., Lorenc, A., et al. (2022). Immune disease risk variants regulate gene expression dynamics during CD4+ T cell activation. Nat. Genet. 54, 817–826. 10.1038/s41588-022-01066-3.

11. Aquino, Y., Bisiaux, A., Li, Z., O’Neill, M., Mendoza-Revilla, J., Merkling, S.H., Kerner, G., Hasan, M., Libri, V., Bondet, V., et al. (2023). Dissecting human population variation in single-cell responses to SARS-CoV-2. Nature 621, 120–128. 10.1038/s41586-023-06422-9.

12. Kumasaka, N., Rostom, R., Huang, N., Polanski, K., Meyer, K.B., Patel, S., Boyd, R., Gomez, C., Barnett, S.N., Panousis, N.I., et al. (2023). Mapping interindividual dynamics of innate immune response at single-cell resolution. Nat. Genet. 55, 1066–1075. 10.1038/s41588-023-01421-y.

13. Cuomo, A.S.E., Seaton, D.D., McCarthy, D.J., Martinez, I., Bonder, M.J., Garcia-Bernardo, J., Amatya, S., Madrigal, P., Isaacson, A., Buettner, F., et al. (2020). Single-cell RNA-sequencing of differentiating iPS cells reveals dynamic genetic effects on gene expression. Nat. Commun. 11, 810. 10.1038/s41467-020-14457-z.

14. Jerber, J., Seaton, D.D., Cuomo, A.S.E., Kumasaka, N., Haldane, J., Steer, J., Patel, M., Pearce, D., Andersson, M., Bonder, M.J., et al. (2021). Population-scale single-cell RNA-seq profiling across dopaminergic neuron differentiation. Nat. Genet. 53, 304–312. 10.1038/s41588-021-00801-6.

15. Arthur, T.D., Nguyen, J.P., Henson, B.A., D’Antonio-Chronowska, A., Jaureguy, J., Silva, N., Arias, A.D., Benaglio, P., Berggren, W.T., Borja, V., et al. (2025). Multiomic QTL mapping reveals phenotypic complexity of GWAS loci and prioritizes putative causal variants. Cell Genomics 5. 10.1016/j.xgen.2025.100775.

16. Nguyen, J.P., Arthur, T.D., Fujita, K., Salgado, B.M., Donovan, M.K.R., Matsui, H., Kim, J.H., D’Antonio-Chronowska, A., D’Antonio, M., and Frazer, K.A. (2023). eQTL mapping in fetal-like pancreatic progenitor cells reveals early developmental insights into diabetes risk. Nat. Commun. 14, 6928. 10.1038/s41467-023-42560-4.

17. Knowles, D.A., Burrows, C.K., Blischak, J.D., Patterson, K.M., Serie, D.J., Norton, N., Ober, C., Pritchard, J.K., and Gilad, Y. (2018). Determining the genetic basis of anthracycline-cardiotoxicity by molecular response QTL mapping in induced cardiomyocytes. eLife 7, e33480. 10.7554/eLife.33480.

18. Zhong, Y., De, T., Mishra, M., Avitia, J., Alarcon, C., and Perera, M.A. (2023). Leveraging drug perturbation to reveal genetic regulators of hepatic gene expression in African Americans. Am. J. Hum. Genet. 110, 58–70. 10.1016/j.ajhg.2022.12.005.

19. Popp, J.M., Rhodes, K., Jangi, R., Li, M., Barr, K., Tayeb, K., Battle, A., and Gilad, Y. Cell type and dynamic state govern genetic regulation of gene expression in heterogeneous differentiating cultures. Cell Genomics. 10.1016/j.xgen.2024.100701.

20. Cortal, A., Martignetti, L., Six, E., and Rausell, A. (2021). Gene signature extraction and cell identity recognition at the single-cell level with Cell-ID. Nat. Biotechnol. 39, 1095–1102. 10.1038/s41587-021-00896-6.

21. Muñoz-Sanjuán, I., and Brivanlou, A.H. (2002). Neural induction, the default model and embryonic stem cells. Nat. Rev. Neurosci. 3, 271–280. 10.1038/nrn786.

22. Murry, C.E., and Keller, G. (2008). Differentiation of Embryonic Stem Cells to Clinically Relevant Populations: Lessons from Embryonic Development. Cell 132, 661–680. 10.1016/j.cell.2008.02.008.

23. Kamiya, D., Banno, S., Sasai, N., Ohgushi, M., Inomata, H., Watanabe, K., Kawada, M., Yakura, R., Kiyonari, H., Nakao, K., et al. (2011). Intrinsic transition of embryonic stem-cell differentiation into neural progenitors. Nature 470, 503–509. 10.1038/nature09726.

24. Li, X., Yue, X., Pastor, W.A., Lin, L., Georges, R., Chavez, L., Evans, S.M., and Rao, A. (2016). Tet proteins influence the balance between neuroectodermal and mesodermal fate choice by inhibiting Wnt signaling. Proc. Natl. Acad. Sci. 113, E8267–E8276. 10.1073/pnas.1617802113.

25. Dann, E., Henderson, N.C., Teichmann, S.A., Morgan, M.D., and Marioni, J.C. (2022). Differential abundance testing on single-cell data using k-nearest neighbor graphs. Nat. Biotechnol. 40, 245–253. 10.1038/s41587-021-01033-z.

26. Chen, S., and Charness, M.E. (2008). Ethanol inhibits neuronal differentiation by disrupting activity-dependent neuroprotective protein signaling. Proc. Natl. Acad. Sci. 105, 19962–19967. 10.1073/pnas.0807758105.

27. Hellmann, J., Rommelspacher, H., and Wernicke, C. (2009). Long-Term Ethanol Exposure Impairs Neuronal Differentiation of Human Neuroblastoma Cells Involving Neurotrophin-Mediated Intracellular Signaling and in Particular Protein Kinase C. Alcohol. Clin. Exp. Res. 33, 538–550. 10.1111/j.1530-0277.2008.00867.x.

28. Guadagnoli, T., Caltana, L., Vacotto, M., Gironacci, M.M., and Brusco, A. (2016). Direct effects of ethanol on neuronal differentiation: An in vitro analysis of viability and morphology. Brain Res. Bull. 127, 177–186. 10.1016/j.brainresbull.2016.09.013.

29. Setty, M., Kiseliovas, V., Levine, J., Gayoso, A., Mazutis, L., and Pe’er, D. (2019). Characterization of cell fate probabilities in single-cell data with Palantir. Nat. Biotechnol. 37, 451–460. 10.1038/s41587-019-0068-4.

30. He, B., Wen, Y., Hu, S., Wang, G., Hu, W., Magdalou, J., Chen, L., and Wang, H. (2019). Prenatal caffeine exposure induces liver developmental dysfunction in offspring rats. J. Endocrinol. 242, 211–226. 10.1530/JOE-19-0066.

31. Goette, A., Lendeckel, U., Kuchenbecker, A., Bukowska, A., Peters, B., Klein, H.U., Huth, C., and Röcken, C. (2007). Cigarette smoking induces atrial fibrosis in humans via nicotine. Heart 93, 1056–1063. 10.1136/hrt.2005.087171.

32. Zhou, X., Sheng, Y., Yang, R., and Kong, X. (2010). Nicotine Promotes Cardiomyocyte Apoptosis via Oxidative Stress and Altered Apoptosis-Related Gene Expression. Cardiology 115, 243–250. 10.1159/000301278.

33. Taylor-Weiner, A., Aguet, F., Haradhvala, N.J., Gosai, S., Anand, S., Kim, J., Ardlie, K., Van Allen, E.M., and Getz, G. (2019). Scaling computational genomics to millions of individuals with GPUs. Genome Biol. 20, 228. 10.1186/s13059-019-1836-7.

34. Koscielny, G., An, P., Carvalho-Silva, D., Cham, J.A., Fumis, L., Gasparyan, R., Hasan, S., Karamanis, N., Maguire, M., Papa, E., et al. (2016). Open Targets: a platform for therapeutic target identification and validation. Nucleic Acids Res. 45, D985. 10.1093/nar/gkw1055.

35. Mostafavi, H., Spence, J.P., Naqvi, S., and Pritchard, J.K. (2023). Systematic differences in discovery of genetic effects on gene expression and complex traits. Nat. Genet. 55, 1866–1875. 10.1038/s41588-023-01529-1.

36. Moore, J.E., Purcaro, M.J., Pratt, H.E., Epstein, C.B., Shoresh, N., Adrian, J., Kawli, T., Davis, C.A., Dobin, A., Kaul, R., et al. (2020). Expanded encyclopaedias of DNA elements in the human and mouse genomes. Nature 583, 699–710. 10.1038/s41586-020-2493-4.

37. Maunakea, A.K., Nagarajan, R.P., Bilenky, M., Ballinger, T.J., D’Souza, C., Fouse, S.D., Johnson, B.E., Hong, C., Nielsen, C., Zhao, Y., et al. (2010). Conserved role of intragenic DNA methylation in regulating alternative promoters. Nature 466, 253–257. 10.1038/nature09165.

38. Forrest, A.R.R., Kawaji, H., Rehli, M., Kenneth Baillie, J., de Hoon, M.J.L., Haberle, V., Lassmann, T., Kulakovskiy, I.V., Lizio, M., Itoh, M., et al. (2014). A promoter-level mammalian expression atlas. Nature 507, 462–470. 10.1038/nature13182.

39. Bonn, S., Zinzen, R.P., Girardot, C., Gustafson, E.H., Perez-Gonzalez, A., Delhomme, N., Ghavi-Helm, Y., Wilczyński, B., Riddell, A., and Furlong, E.E.M. (2012). Tissue-specific analysis of chromatin state identifies temporal signatures of enhancer activity during embryonic development. Nat. Genet. 44, 148–156. 10.1038/ng.1064.

40. Arner, E., Daub, C.O., Vitting-Seerup, K., Andersson, R., Lilje, B., Drabløs, F., Lennartsson, A., Rönnerblad, M., Hrydziuszko, O., Vitezic, M., et al. (2015). Transcribed enhancers lead waves of coordinated transcription in transitioning mammalian cells. Science. 10.1126/science.1259418.

41. Anderson, S.T., and FitzGerald, G.A. (2020). Sexual dimorphism in body clocks. Science 369, 1164–1165. 10.1126/science.abd4964.

42. Nasser, J., Bergman, D.T., Fulco, C.P., Guckelberger, P., Doughty, B.R., Patwardhan, T.A., Jones, T.R., Nguyen, T.H., Ulirsch, J.C., Lekschas, F., et al. (2021). Genome-wide enhancer maps link risk variants to disease genes. Nature 593, 238–243. 10.1038/s41586-021-03446-x.

43. Exome Aggregation Consortium, Lek, M., Karczewski, K.J., Minikel, E.V., Samocha, K.E., Banks, E., Fennell, T., O’Donnell-Luria, A.H., Ware, J.S., Hill, A.J., et al. (2016). Analysis of protein-coding genetic variation in 60,706 humans. Nature 536, 285–291. 10.1038/nature19057.

44. Sakaue, S., Kanai, M., Tanigawa, Y., Karjalainen, J., Kurki, M., Koshiba, S., Narita, A., Konuma, T., Yamamoto, K., Akiyama, M., et al. (2021). A cross-population atlas of genetic associations for 220 human phenotypes. Nat. Genet. 53, 1415–1424. 10.1038/s41588-021-00931-x.

45. Fair, B., Buen Abad Najar, C.F., Zhao, J., Lozano, S., Reilly, A., Mossian, G., Staley, J.P., Wang, J., and Li, Y.I. (2024). Global impact of unproductive splicing on human gene expression. Nat. Genet. 56, 1851–1861. 10.1038/s41588-024-01872-x.

46. Foley, C.N., Staley, J.R., Breen, P.G., Sun, B.B., Kirk, P.D.W., Burgess, S., and Howson, J.M.M. (2021). A fast and efficient colocalization algorithm for identifying shared genetic risk factors across multiple traits. Nat. Commun. 12, 764. 10.1038/s41467-020-20885-8.

47. Nicolae, D.L., Gamazon, E., Zhang, W., Duan, S., Dolan, M.E., and Cox, N.J. (2010). Trait-Associated SNPs Are More Likely to Be eQTLs: Annotation to Enhance Discovery from GWAS. PLOS Genet. 6, e1000888. 10.1371/journal.pgen.1000888.

48. de Lange, K.M., Moutsianas, L., Lee, J.C., Lamb, C.A., Luo, Y., Kennedy, N.A., Jostins, L., Rice, D.L., Gutierrez-Achury, J., Ji, S.-G., et al. (2017). Genome-wide association study implicates immune activation of multiple integrin genes in inflammatory bowel disease. Nat. Genet. 49, 256–261. 10.1038/ng.3760.

49. Thakurela, S., Sahu, S.K., Garding, A., and Tiwari, V.K. (2015). Dynamics and function of distal regulatory elements during neurogenesis and neuroplasticity. Genome Res. 25, 1309–1324. 10.1101/gr.190926.115.

50. Bravo González-Blas, C., Quan, X., Duran-Romaña, R., Taskiran, I.I., Koldere, D., Davie, K., Christiaens, V., Makhzami, S., Hulselmans, G., de Waegeneer, M., et al. (2020). Identification of genomic enhancers through spatial integration of single-cell transcriptomics and epigenomics. Mol. Syst. Biol. 16, e9438. 10.15252/msb.20209438.

51. Mitschka, S., and Mayr, C. (2022). Context-specific regulation and function of mRNA alternative polyadenylation. Nat. Rev. Mol. Cell Biol. 23, 779–796. 10.1038/s41580-022-00507-5.

52. Lin, W., Wall, J.D., Li, G., Newman, D., Yang, Y., Abney, M., VandeBerg, J.L., Olivier, M., Gilad, Y., and Cox, L.A. (2024). Genetic regulatory effects in response to a high-cholesterol, high-fat diet in baboons. Cell Genomics 4. 10.1016/j.xgen.2024.100509.

53. Huang, Y., McCarthy, D.J., and Stegle, O. (2019). Vireo: Bayesian demultiplexing of pooled single-cell RNA-seq data without genotype reference. Genome Biol. 20, 273. 10.1186/s13059-019-1865-2.

54. Byrska-Bishop, M., Evani, U.S., Zhao, X., Basile, A.O., Abel, H.J., Regier, A.A., Corvelo, A., Clarke, W.E., Musunuri, R., Nagulapalli, K., et al. (2022). High-coverage whole-genome sequencing of the expanded 1000 Genomes Project cohort including 602 trios. Cell 185, 3426–3440.e19. 10.1016/j.cell.2022.08.004.

55. Cao, J., O’Day, D.R., Pliner, H.A., Kingsley, P.D., Deng, M., Daza, R.M., Zager, M.A., Aldinger, K.A., Blecher-Gonen, R., Zhang, F., et al. (2020). A human cell atlas of fetal gene expression. Science 370, eaba7721. 10.1126/science.aba7721.

56. Braun, E., Danan-Gotthold, M., Borm, L.E., Lee, K.W., Vinsland, E., Lönnerberg, P., Hu, L., Li, X., He, X., Andrusivová, Ž., et al. (2023). Comprehensive cell atlas of the first-trimester developing human brain. Science 382, eadf1226. 10.1126/science.adf1226.

57. Hoffman, G.E., Lee, D., Bendl, J., Prashant, N.M., Hong, A., Casey, C., Alvia, M., Shao, Z., Argyriou, S., Therrien, K., et al. (2024). Efficient differential expression analysis of large-scale single cell transcriptomics data using dreamlet. BioRxiv Prepr. Serv. Biol., 2023.03.17.533005. 10.1101/2023.03.17.533005.

58. Korotkevich, G., Sukhov, V., Budin, N., Shpak, B., Artyomov, M.N., and Sergushichev, A. (2021). Fast gene set enrichment analysis. Preprint at bioRxiv, 10.1101/060012 https://doi.org/10.1101/060012.

59. Liberzon, A., Birger, C., Thorvaldsdóttir, H., Ghandi, M., Mesirov, J.P., and Tamayo, P. (2015). The Molecular Signatures Database Hallmark Gene Set Collection. Cell Syst. 1, 417–425. 10.1016/j.cels.2015.12.004.

60. Weinberger, E., Lin, C., and Lee, S.-I. (2023). Isolating salient variations of interest in single-cell data with contrastiveVI. Nat. Methods 20, 1336–1345. 10.1038/s41592-023-01955-3.

61. Wolf, F.A., Angerer, P., and Theis, F.J. (2018). SCANPY: large-scale single-cell gene expression data analysis. Genome Biol. 19, 15. 10.1186/s13059-017-1382-0.

62. Shabalin, A.A. (2012). Matrix eQTL: ultra fast eQTL analysis via large matrix operations. Bioinformatics 28, 1353–1358. 10.1093/bioinformatics/bts163.

63. Davis, J.R., Fresard, L., Knowles, D.A., Pala, M., Bustamante, C.D., Battle, A., and Montgomery, S.B. (2016). An Efficient Multiple-Testing Adjustment for eQTL Studies that Accounts for Linkage Disequilibrium between Variants. Am. J. Hum. Genet. 98, 216–224. 10.1016/j.ajhg.2015.11.021.

64. Urbut, S.M., Wang, G., Carbonetto, P., and Stephens, M. (2019). Flexible statistical methods for estimating and testing effects in genomic studies with multiple conditions. Nat. Genet. 51, 187–195. 10.1038/s41588-018-0268-8.

65. Elorbany, R., Popp, J.M., Rhodes, K., Strober, B.J., Barr, K., Qi, G., Gilad, Y., and Battle, A. (2022). Single-cell sequencing reveals lineage-specific dynamic genetic regulation of gene expression during human cardiomyocyte differentiation. PLOS Genet. 18, e1009666. 10.1371/journal.pgen.1009666.

66. Wold, H. (1975). Path Models with Latent Variables: The NIPALS Approach*. In Quantitative Sociology International Perspectives on Mathematical and Statistical Modeling., H. M. Blalock, A. Aganbegian, F. M. Borodkin, R. Boudon, and V. Capecchi, eds. (Academic Press), pp. 307–357. 10.1016/B978-0-12-103950-9.50017-4.

67. Chen, S., Francioli, L.C., Goodrich, J.K., Collins, R.L., Kanai, M., Wang, Q., Alföldi, J., Watts, N.A., Vittal, C., Gauthier, L.D., et al. (2024). A genomic mutational constraint map using variation in 76,156 human genomes. Nature 625, 92–100. 10.1038/s41586-023-06045-0.

68. Lambert, S.A., Jolma, A., Campitelli, L.F., Das, P.K., Yin, Y., Albu, M., Chen, X., Taipale, J., Hughes, T.R., and Weirauch, M.T. (2018). The Human Transcription Factors. Cell 172, 650–665. 10.1016/j.cell.2018.01.029.

